# Ecological niche models as hypothesis generators of functional genetic differentiation and potential local adaptation in a Mediterranean alpine ecosystem

**DOI:** 10.1101/2020.02.18.954867

**Authors:** Javier Morente-López, Jamie M. Kass, Carlos Lara-Romero, Josep María Serra-Diaz, José Carmen Soto-Correa, Robert P. Anderson, José María Iriondo

## Abstract

Geographically disparate populations within a species’ range may show important differences including variation in ecological, demographic, genetic and phenotypic characteristics. Based on the Center-Periphery Hypothesis, it is often assumed that environmental conditions are optimal in the geographic center of the range and stressful or suboptimal at the periphery, implying ecological marginality is concordant with geographic periphery. But this assumption has been challenged as geographical and ecological gradients are not necessarily concordant. The conservation value of populations inhabiting environmentally marginal areas is still under debate and is closely related with their evolutionary potential. Strong selective pressures caused by stressful conditions may generate novel adaptations in marginal areas, conferring these populations distinct evolutionary potential. But populations inhabiting marginal areas may also show reductions in neutral and adaptive genetic diversity via drift and inbreeding.

In this work we explore the potential of ecological niche models (ENMs) to identify environmentally optimal and marginal areas, as well as the principal putative selective pressures likely to act. To do so, we built a carefully parameterized ENM of *Silene ciliata*, a dominant plant species of Mediterranean alpine habitats. Complementarily, we selected wild populations inhabiting contrasting environmental conditions and carried out common garden experiments to detect genetic differentiation among populations associated with functional traits. With the resulting information, we tested whether environmentally marginal populations defined by the ENM had genetically differentiated phenotypes that are potentially adaptive and, thus, of conservation value.

We found genetically based phenotypic differentiation of phenological traits between populations inhabiting areas identified by the ENM as marginal and optimal, as well as between populations with different habitat suitability values. Results supported ENMs as powerful tools for determining environmental marginality and identifying selection pressures, and thus also as hypothesis generators for divergent selection. Furthermore, genetically based phenotypic differentiation found underlines the potential adaptive value of populations inhabiting marginal areas. The approach developed here provides a theoretically justified and practical way to study adaptive processes and provide insights about the conservation value of marginal populations.

## 1. Introduction

Understanding key aspects of species distributions has long been a major concern in biology (Brown, 1984; Grinnell, 1917; Hutchinson, 1957). At the heart of theoretical relationships between niche and distribution, the Center-Periphery Hypothesis (hereafter CPH) posits that geographically disparate populations within a species’ range may have important differences (Abeli et al., 2014; Soule, 1973), which can include variation in demographic, genetic and phenotypic characteristics (Gaston and Sheffield, 2009; Pironon et al., 2017; Sexton et al., 2009). Understanding the potential effects of such variation is a topic of great interest in ecology, biogeography, evolution, and conservation biology (Abeli et al., 2014; Abeli and Orsenigo, 2018; Brussard, 1984; Eckert et al., 2008; Gaston and Sheffield, 2009; Holt and Keitt, 2005; Kawecki, 2008; Lesica and Allendorf, 1995; Pouget et al., 2013; Sexton et al., 2009).

Different factors have been considered to classify an area as central or peripheral (Pironon et al., 2017). It is often assumed that environmental conditions are optimal in the geographic center of a species’ range and impoverished or suboptimal at the periphery (Brown, 1984; Hoffmann and Blows, 1994; Holt and Keitt, 2005), which can lead to the assumption that peripheral populations are ecologically marginal. Hence, it has been proposed that peripheral populations occupy smaller and more fragmented areas, exhibit decreased abundance, declining demographic trends, and low individual fitness (Angert, 2006; Brown, 1984; Hengeveld and Haeck, 1982; Herlihy and Eckert, 2005; Villellas et al., 2013). However, this assumption has been challenged since geographical and ecological gradients are not necessarily concordant (Hardie and Hutchings, 2010; Pironon et al., 2015; Soule, 1973) and the corresponding abundance, demographic and fitness patterns found are not always consistent (Abeli et al., 2014; Pironon et al., 2017).

Opposing viewpoints have also been presented regarding the genetic patterns and properties of geographically and/or environmentally marginal populations (Eckert et al., 2008; Kawecki, 2008). On the one hand, the demographic impoverishment underlying the classical formulation of the CPH would lead to reduced effective population size and lower gene flow among populations, and consequently, reductions in the neutral and adaptive genetic diversity levels through genetic drift and inbreeding processes (Eckert et al., 2008; Kawecki, 2008). As a result, marginal populations may end up with a significant genetic load, maladapted individuals due to loss of heterosis and deleterious alleles accumulation. On the other hand, strong selection pressures caused by stressful conditions may generate novel adaptations in marginal areas, conferring these populations distinct evolutionary potential (Kawecki, 2008; Rolland et al., 2015) and therefore adaptive value. In this context of genetic impoverishment vs. adaptive potential, the conservation value of populations inhabiting environmentally marginal areas is a long-standing and debate in ecology (Abeli et al., 2014; Abeli and Orsenigo, 2018; Channell, 2004; Hunter and Hutchinson, 1994; Lesica and Allendorf, 1995; Millar and Libby, 1991).

Identifying genetically based phenotypic variation is a key prerequisite for assessing local adaptation processes related to contrasting environmental conditions within species’ ranges and constitutes a key issue in evolutionary and conservation biology. Some types of functional traits may respond to differing environments with different sensitivities and thus play a more important role in divergent selection and adaptation. Accordingly, life-history traits and other closely related traits like phenology have been suggested to be more likely affected by adaptation to climate change than physiological traits (Bradshaw and Holzapfel, 2008; Chuine, 2010; Hoffmann and Sgró, 2011). However, physiological traits representing the species’ capacity to deal with extreme environmental pressures (such as high and low temperatures cell resistance) are also of great interest in this context, especially in ecosystems where extreme temperatures occur (Nobel et al., 2002). The study of variation in different types of functional traits can help identify patterns of environmental marginality and the environmental factors responsible for divergent selection and potential local adaptation.

Ecological niche models (hereafter ENMs) can be a powerful correlative tool to detect environmentally marginal areas within a species’ range and help generate hypotheses regarding the most important limiting environmental variables. ENMs estimate habitat suitability values (hereafter HS) based on modeled responses of the species to its environment (Anderson and Raza, 2010; Phillips and Dudík, 2008). ENMs are widely used to study the climatic drivers of species distributions and their shifts caused by climate change (Guisan and Thuiller, 2005; Peterson et al., 2011). They usually assume no functional genetic variation across the focal species’ range for factors related to the species’ distributional ecology, and thus also to functional responses to the environmental variables used as model predictors (Anderson, 2013; Hällfors et al., 2016). But an adaptive response can occur *in situ* if there are contrasting selective pressures within the species’ range and enough variation in genes responsible for local adaptation (Blanquart et al., 2013; Kawecki and Ebert, 2004). Mechanistic models could have more realistic predictions than correlative ENMs regarding the fine-scale ecology of the species (Kearney and Porter, 2009). But great time and resources are consumed in field studies to obtain the variables required to parameterize mechanistic models, such as dispersal, competition-facilitation relationships, or physiological responses of species (Benito-Garzón et al., 2011; Kearney and Porter, 2009). Although ENMs are correlative models that do not account for the biological processes underlying genetic differentiation and adaptation of a species to its environment (Hällfors et al., 2016; Valladares et al., 2014), they can identify areas with different levels of environmental suitability for the species within its range. In this context, correlative models may be useful hypothesis generators (Dixon and Busch, 2017) and may serve as an initial step to address fine-scale aspects of the species ecology that then could be tested using experimental data. Thus, using ENMs may be possible not only to estimate the distribution of a species and to identify putative important environmental variables that shape it (Searcy and Shaffer, 2016), but also to hypothesize intra-specific and genetically based phenotypic variability mediated by divergent selection within the species’ range. This can be very useful in the context of studies that focus on selection-mediated genetic differentiation and local adaptation. In this context, there is a need for more ecologically realistic niche models (Guevara et al., 2018; Moritz and Agudo, 2013; Owens et al., 2013), especially for species with suspected local adaptation. This approach can be used to identify populations likely to show functional genetic differentiation and thus guide studies that test for this and assess the conservation value of those populations. It is important to consider that model development needs careful consideration closely related with modeling objectives in order to achieve desired performance (Araújo et al., 2019).

The purpose of this work was to explore the value of ENMs as a tool to identify environmentally optimal and marginal areas and to test for genetically based phenotypic differentiation of populations inhabiting marginal areas. As alpine ecosystems are highly variable at a local scale (i.e., small changes in elevation can cause large changes in temperature, humidity, exposure and other variables) (Körner, 2003), selective pressures can change greatly over short distances (Herrera and Bazaga, 2008). Therefore, we focused on *Silene ciliata*, a Mediterranean alpine cushion plant that is an suitable system for the evaluation of evolutionary responses to ongoing warming via climate change in Mediterranean alpine ecosystems (e.g. García-Fernández et al., 2012b, 2013, Giménez-Benavides et al., 2011b, 2018). We used ENMs as hypothesis generators to identify optimal and marginal areas and to guide tests of the association between habitat suitability and functional traits of adaptive value. We specifically addressed three main questions: (i) Can ENMs be sensitive enough to identify contrasting selective pressures inside species distributions, even for species that respond to the environment over small geographic distances? (ii) Can ENM habitat suitability values be used as effective predictors of populations that are genetically differentiated by natural selection? and (iii) Which type of functional traits (i.e., phenological vs. physiological traits) is more likely to respond to differences in habitat suitability values and thus to diverging selective pressures?

To address these questions, we first built an ENM to identify important environmental variables associated with the distribution of the species (and differences in environmental suitability across its range). Then, we carried out common garden experiments to detect genetic differentiation among populations associated with functional traits. With the resulting information, we tested whether environmentally marginal populations had genetically differentiated phenotypes that are potentially adaptive and, thus, of conservation value.

## 2. Materials and Methods

### 2.1 Study system. Study species

*Silene ciliata* Poiret (Caryophyllaceae) is a dwarf cushion perennial plant that constitutes one of the dominant species of Mediterranean alpine habitats. The flowering period spans from the end of July until the end of August. Seeds are relatively small (mean ± SD: 1.5 ± 0.5 mm diameter), and most of them need low-temperature conditions to break seed dormancy and germinate (cold stratification) (Baskin and Baskin, 1998; Giménez-Benavides et al., 2005). Effective pollen and seed dispersal distances are low and relatively invariant across populations (Lara-Romero et al., 2014b, 2016b).

The distribution of *S. ciliata* comprises the mountain ranges of the Northern Mediterranean area from Spain to Bulgaria (see Tutin et al., 1964; Kyrkou et al., 2015), reaching its southernmost limit in the Sistema Central of the Iberian Peninsula (SC). The SC is a southwest-northeast oriented mountain range of approximately 500 km, located in the center of the Iberian Peninsula and composed by three main mountain ranges: Béjar (BJR), Gredos (GRD) and Guadarrama (GDM). *Silene ciliata* populations from the SC have the same phylogenetic origin (demographic history) as shown by chloroplast DNA analysis (Kyrkou et al., 2015). Similar levels of genetic diversity and low genetic differentiation were found among *S. ciliata* populations located along an elevational gradient in the SC, indicating the existence of substantial gene flow across different elevations (García-Fernández et al., 2012; Lara-Romero et al., 2016, Morente-López et al., 2018). In these areas the species grows from 1850 m to the highest mountain altitudes (2592 m) in pastures above the tree line thriving under dry conditions and low temperatures. This Mediterranean Alpine ecosystem presents marked environmental gradients characterized by a strong daily and seasonal temperature fluctuation, a long period of snow cover and a pronounced summer drought combined with high solar radiation (Rivas-Martínez et al., 1990). Vegetation composition of these habitats varies consistently with elevation but is otherwise rather homogeneous throughout the SC (Escudero, Giménez-Benavides, Iriondo, & Rubio, 2004).

### 2.2 Species distribution model construction and evaluation

#### Input data

We used species occurrence data in the SC obtained from the Global Biodiversity Information Facility (GBIF; accessed October 2013) and complemented this dataset with ten new records from us with specimens deposited in the Real Jardín Botánico de Madrid herbarium (Appendix 1). *Silene ciliata* is easily identifiable from other sympatric species of the same genus in the alpine areas of the SC (*Silene boryi* Boiss., which inhabits crag environments). Occurrence records with less than 10 meters accuracy in geographical coordinates were removed. After cleaning the data, a total of 120 presence records were used for model training. To account for sampling biases (Boria et al., 2014; Merow et al., 2014) and subsequent artifactual spatial autocorrelation and environmental biases (Radosavljevic and Anderson, 2014), we spatially thinned our occurrence records using the R package *spThin* 0.1.0.1 (Aiello□Lammens et al., 2015). We selected a thinning distance that ensures no two records were within 0.75 km of one another, yielding 87 records in the final dataset. This relatively low thinning distance was chosen due to the high environmental heterogeneity of the mountainous regions characterizing our study region (Boria et al., 2014). In this context, it was essential to limit the loss of observed environmental variation (by avoiding thinning distances that are too high) (Anderson, 2012), but also to address sampling bias (by avoiding thinning distances that are too low) (Appendix 2).

As our aim for modeling was explanatory rather than predictive (i.e., identifying putative driving variables and environmentally differentiated habitats rather than merely predicting suitability) (Araújo et al., 2019; Araujo and Guisan, 2006; Elith and Leathwick, 2009; Merow et al., 2014), we designed a process to identify and select variables that we hypothesized could explain this species’ distributional patterns and further characterize intra-range environmental variation and extreme environmental conditions (Appendix 3). As a result, we selected the following variables: minimum temperature of the coldest month, precipitation of the driest and the wettest month, mean snowpack calculated in thaw months (February, March and April; López-Moreno et al., 2007) and mean annual potential evapotranspiration. All variables have a 200 m resolution and were obtained from the climatic digital atlas of the Iberian Peninsula (Ninyerola et al., 2005, accessed October 2016). All variables’ pairwise correlation values were below 0.7 (see Appendix 3 for detailed information).

#### Ecological niche model parameterization, evaluation, and selection

We used the ENM algorithm Maxent 3.3.3 (Phillips et al., 2006), a presence-background machine learning technique. As proper background area selection is crucial (Saupe et al., 2012), we focused on defining one that includes the present distribution of the species and a wide variety of environments within the study region. Specifically, we defined a background area by a 10 km buffer around all presence data (after cleaning but before thinning) that excludes areas below 1000 m elevation where the species is not likely to be present. This area includes the environmental heterogeneity accessible to the species considering dispersal limitations and biogeographical barriers conformation (Anderson and Raza, 2010).

To select the model settings with an optimal level of complexity for realistic hypotheses regarding the environmental variables driving the species’ range (Araujo and Guisan, 2006; Galante et al., 2017; Merow et al., 2014), we built multiple models with different settings to compare between them. We considered four combinations of feature classes (FC: Linear, Linear-Quadratic, Hinge and Linear-Quadratic-Hinge), which allow for varying levels of model complexity, and regularization multipliers (RM) (1 through 5 by intervals of 0.5), which penalize complexity as they increase (Phillips and Dudík, 2008). We first selected model settings that resulted in the lowest Akaike Information Criterion corrected for small sample sizes (AIC_C_). We then evaluated selected models by calculating cross-validation summary statistics using a spatially segregated *k*-fold data checkerboard partition method (“checkerboard2” from ENMeval; aggregation factor 2) (Muscarella et al., 2014). Threshold-dependent (omission rate or OR10, using a threshold set by 10% training omission rate) and threshold-independent evaluation statistics (AUC_TEST_, AUC for testing points) were used to evaluate the models in terms of over-fitting and discrimination (see Anderson and Gonzalez, 2011). Model evaluation was performed using the R package *ENMeval* 0.3.0 (Muscarella et al., 2014). In order to consider the full representation of environmental variability inside the species’ distribution, we included all pixels within the study area as background samples in the model (Guevara et al., 2018).

#### Final model and environmental variables importance

We projected the optimal model to the entire study area to predict HS (Figure 1), which we used to distinguish the degree of environmental marginality of different areas inside the range of *S. ciliata*. We classified the study area into “optimal” and “marginal” categories based on the predicted HS values for the spatially thinned localities. The optimal category was arbitrarily assigned to areas with habitat suitability values in the highest 33^rd^ percentile of the HS value distribution, whereas the marginal category was assigned to those in the lowest 33^rd^ percentile. The habitat suitability values comprised between the lowest and the highest 33^rd^ percentile correspond to those which are around the median and were not used in our analyses since we wanted to focus on contrasting environmental conditions.

**Figure 1:**
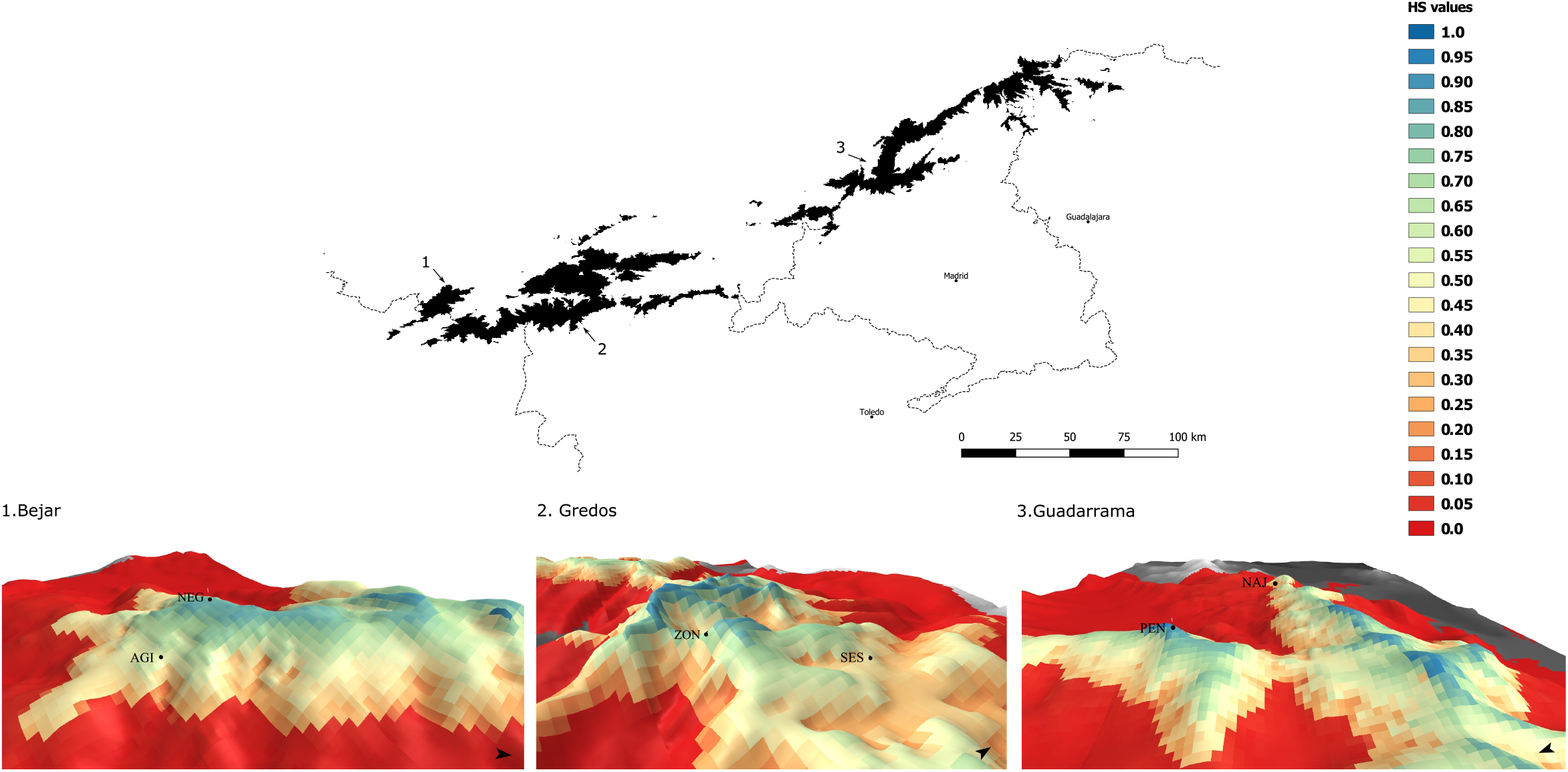
*Silene ciliata* ecological niche model (ENM) in the Sistema Central of the Iberian Peninsula. Black areas represent elevations above 1500m. Each of the maps at the bottom represents the geographical projection of the habitat suitability values obtained with the selected ENM in each of the areas of the Sistema Central where the phenotypic differentiation study was developed (squared areas), shown over elevation. The six populations used for phenotypic study are represented in the model maps (Table 1).

Based on the permutation importance and also regarding at parameter weights (lambda values) from the ENM (see Phillips, 2011; Searcy and Shaffer, 2016 for a detailed definition of permutation importance), we selected the environmental variables that contribute most to explain the *S. ciliata* modeled HS values (Galante et al., 2017; Searcy and Shaffer, 2016). We plotted the environmental variable response curves in order to study the direction of the environmental drivers that potentially generate the main selective pressures that make areas environmentally marginal. We also plotted the environmental coordinates of the presence data in the environmental space of the most important variables with non-zero coefficients in the model (see Appendix 4). We examined where the presence points belonging to the populations selected for the phenotypic differentiation study were located in the environmental space. We also examined which mountain each of these presence records originated from to determine if there is any environmental differentiation between mountains within the SC.

**Table 1:**
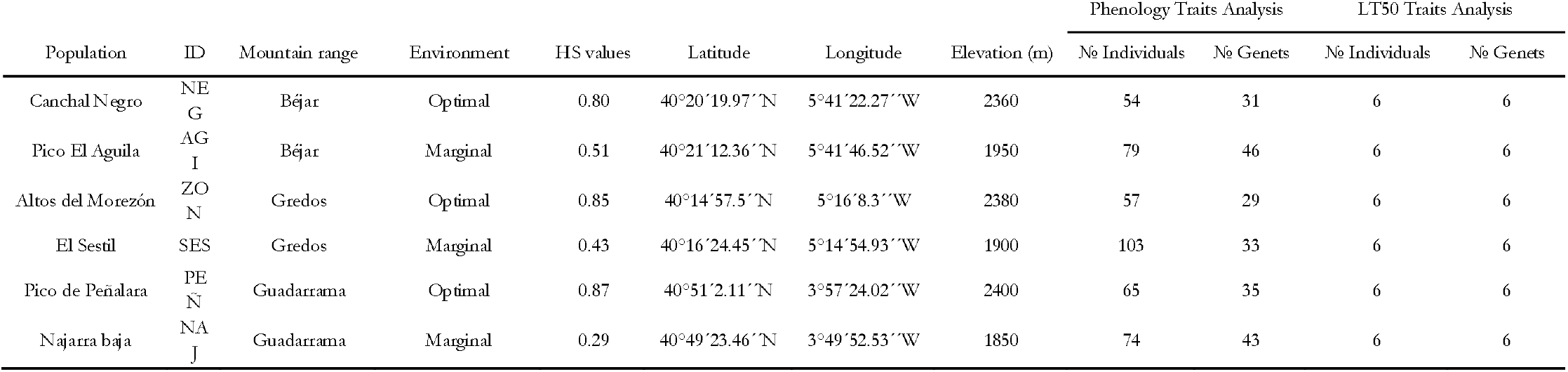
*Silene ciliata* populations of the Sistema Central selected for phenotypic differentiation study. For each population its name and ID, mountain range of origin, environmental classification, habitat suitability values (HS) extracted from the Maxent model and geographical coordinates are provided. Also the sample size (number of individuals and genets) for the phenology and lethal temperature (LT50) phenotypic measurements is provided. Environmental classification in optimal and marginal areas was made based on HS values (see methods).

### 2.3 Genetically-based phenotypic differentiation analysis

#### Common garden experiments and functional traits measurements

##### Flowering phenology

In each of the three mountains ranges of the SC, we selected two populations in the field located in areas defined as “optimal” (high HS) and “marginal” (low HS but still sufficient to support reproducing populations) (Fig 1 and Table 1). At the end of the summer of 2013 we collected a minimum of thirty randomly selected plants from these six selected populations. The collected plants were divided into a variable number of cuttings (depending on the size of the plant) to obtain a greater number of genetically identical individuals (genets) to overcome future losses during rooting and growth. *Silene ciliata* is a slow-growing species that requires at least two years to reach reproductive adulthood from seeds. Thus, collecting cuttings of adult individuals is an effective way to quickly obtain reproductive individuals. The whole collection of cuttings was allowed to root and grow in the Universidad Rey Juan Carlos CULTIVE laboratory (690 m a.s.l.) under common garden conditions for seven months to minimize carry-over effects from the original environment. In March 2014, a total of 2200 plants were transplanted into 2.5 L pots. Data on flowering phenology were collected in the growing season of 2014 from two cuttings of each genet when possible, yielding a minimum of 54 individuals and 29 genets from each population. The diameter of all individuals was measured on the date when the first plant started flowering (15 April 2014). Since then and until 25 September 2014, when the last new flower appeared, we counted the total number of new flowers of each plant weekly with a total of 23 censuses. From these data we calculated five individual-based flowering traits: flowering onset, peak, end and duration, and total number of flowers. Flowering onset and end were defined as the number of days elapsed between 1 January 2014 and the dates in which the first and last flower of each plant appeared, respectively. Duration was estimated as the number of days elapsed between the onset and end of flowering. Peak was defined as the number of days elapsed between the date of flowering onset of an individual and the date with the greatest number of open flowers on that individual.

##### Cell resistance to extreme temperatures

At the end of the summer of 2013 we collected seeds from a minimum of thirty randomly selected plants from the six selected populations. Seeds were germinated and seedlings were grown for eight months before data collection under common garden conditions in the Universidad Rey Juan Carlos CULTIVE laboratory. Tolerance to extreme temperatures was determined in May-Jun 2014 by measuring electrolyte leakage, an indicator of cell membrane integrity (Didden-Zopfy and Nobel, 1982; Drennan, 2009; Soto-Correa et al., 2013). Six plants per population were randomly selected for the analysis. Six leaf discs of each plant were obtained with a core borer of 2 mm in diameter and placed in 1.5-ml microtubes containing damp cotton to prevent desiccation. Each leaf disc was incubated at a given temperature for 1 h, after which the disc was removed from the microtube, placed in a glass vial containing 15 ml deionized distilled water and shaken at 200 rpm for 40 min. Electrical conductivity was measured with an Orion-3 conductivity meter (Thermo Electron Corp., Marietta, OH, USA). The process was repeated for the following temperature sequences: 5, 0, −5, −10, −15 and −20 °C for low temperatures and 30, 35, 40, 45, 50 and 55 °C for high temperatures. The temperature at which half of the maximum electrolyte leakage was reached was estimated for each plant separately for both low (hereafter LT50_LOW_) and high (LT50_HIGH_) incubation temperature sequences (see Nobel et al., 2002 detailed description). LT50 values have been found to be good predictors of extreme temperature tolerance in the field (Drennan, 2009; Nobel et al., 2002).

#### Data analysis: Categorical approach

We analyzed whether our defined optimal (high HS) and marginal (low HS) population categories, identified through ENMs, was a determining factor of the recorded functional traits using two different statistical analyses.

For *flowering phenology* we used Cox proportional hazards regression models. These models are used to analyze time-to-event data such as *flowering phenology*. We used the function *coxme* from the R package *coxme* version 2.2–7 (Therneau, 2018) for mixed effects Cox models. We modeled rate of flowering over time with plant size as a covariate, environmental classes as a fixed factor with two levels (optimal/marginal), and population of origin as a random factor.

In the case of *cell resistance to extreme temperatures*, we tested the association between physiological traits (LT50_HIGH_ and LT50_LOW_) and environmental classes using linear mixed models (LMMs) with the same model design explained above for phenological traits. We used the function *lmer* from the R package *lme4* version 1.1–17 (Bates et al., 2014) to perform the analysis.

#### Data analysis: Continuous approach

To assess whether HS values can be used as predictors of genetically-based phenotypic differentiation, we used linear models. We assessed (1) *flowering phenology* (onset, peak, duration and end of flowering) and (2) *cell resistance to extreme temperatures* (LT50_HIGH_ and LT50_LOW_) to continuous HS values and plant size.

In all analyses we calculated Wald *χ*^2^ tests for each fixed effect of the models explained above. *P*-values were estimated using function *Anova* from R package *car* version 2.1-4 (Fox & Weisberg, 2011). Variable significance was calculated using ANOVA type III to deal with the collinearity between variables inherent to our experimental design. Plant size was not considered in the LT50 analysis since data were obtained using artificially cut leaf discs and we assumed no effect of seedling size on this variable.

## 3. Results

### 3.1 Model selection, evaluation and spatial projection

Four models were identified as optimal as they all had delta AIC_c_ values less than two. These four models also had similar values for other evaluation metrics (Mean OR10 and AUC test), had relatively high regulation multiplier values (3 - 5), and had the same feature class combination (LQH; Table 2). The number of model parameters (non-zero coefficients after regularization) for these models ranged from 5 to 7 (Table 2). We selected the model with the lowest AIC_c_ value, which was also one of the simplest with 5 parameters, for further analysis.

**Table 2.** Evaluation metrics of ENMs developed for *S. ciliata* in the Sistema Central of the Península Ibérica. The four models with ΔAIC values below 2 are shown.

| Model | Feature Classes | Regularization Multiplier | $\Delta AIC_c$ | AUC test | Mean Omission Rate 10 | Model Parameters |
| --- | --- | --- | --- | --- | --- | --- |
| 1 | LQH | 4,5 | 0,00 | 0,95 | 0,12 | 5 |
| 2 | LQH | 5 | 0,61 | 0,95 | 0,12 | 5 |
| 3 | LQH | 3 | 1,34 | 0,95 | 0,12 | 7 |
| 4 | LQH | 4 | 1,47 | 0,95 | 0,12 | 6 |

Feature Classes (H = Hinge, L = Linear, Q = Quadratic); $\Delta AIC_c$ , increase of AIC values between models; AUC test, threshold-independent metric based on predicted values for the test localities; Mean Omission Rate 10, threshold-dependent metric that indicates the proportion of test localities with suitability values lower than that which excludes the 10% of training localities with the lowest predicted suitability; Model Parameters, number of parameters in the model.

When we projected this ENM to the background spatial extent, the highest HS values were concentrated in the mountain summits around 2400 m, and lower elevations corresponded with decreasing HS values that were <0.1 below 1800 m (Figure 1). Populations selected for the functional traits differentiation analysis were located in different areas inside the environmental space created with the variables selected by the model (Appendix 4, Figure A 4.1a). No major environmental differentiation was found between mountain ranges (Appendix 4, Figure A 4.1b).

### 3.2 Environmental variables used and response curves

The variable with the highest permutation importance value was snowpack, followed by minimum temperature of the coldest month, potential evapotranspiration and, finally, precipitation of the driest month (Table 3). Precipitation of the wettest month was not considered important by the model, as it had zero permutation importance value and a lambda value of zero. The model response shapes were varied, with two linear (snowpack and minimum temperature of the coldest month), two quadratic (precipitation of the driest month and potential evapotranspiration) and one hinge (snowpack; Table 4). Response curves of each variable showed the type of relation between the respective environmental variable and HS values (Figure 2). As minimum temperature of the coldest month increases, HS values decrease with a clear threshold around −8 °C, at which HS values decrease rapidly, reaching values close to zero at around 3 °C (Figure 2A). HS values are close to zero until snowpack values are positive (presence of snow cover), reaching a maximum when snowpack values reach 1 (Figure 2B). Precipitation of the driest month and potential evapotranspiration showed similar responses (Figure 2C and 2D), with gradually decreasing HS values as both variables increased.

**Table 3.** Variables ordered by the permutation importance obtained from the ENM.

| Variable | Permutation Importance |
| --- | --- |
| Snow Pack | 76,89 |
| BIO6 | 10,32 |
| PET | 7,19 |
| BIO14 | 5,60 |
| BIO 13 | 0 |

Snow Pack, mean snow accumulation ratio in the thaw months; BIO6, min temperature of coldest month; PET, annual potential evapotranspiration; BIO14, precipitation of driest month; BIO13, precipitation of wettest month.

**Table 4.** Feature classes used and lambda values from the ENM model. Variables with zero lambda values were excluded.

| Variable | Distribution | Lambda Value Coefficient |
| --- | --- | --- |
| Snow Pack | Linear | 0,88 |
| Snow Pack | Hinge | -2,48 |
| BIO14 | Quadratic | -2,84 |
| PET | Quadratic | -3,27 |
| BIO6 | Linear | -4,52 |

Snow Pack, mean snow accumulation ratio in the thaw months; PET, annual potential evapotranspiration; BIO14, precipitation of driest month; BIO6, minimum temperature of coldest month.

**Figure 2.**
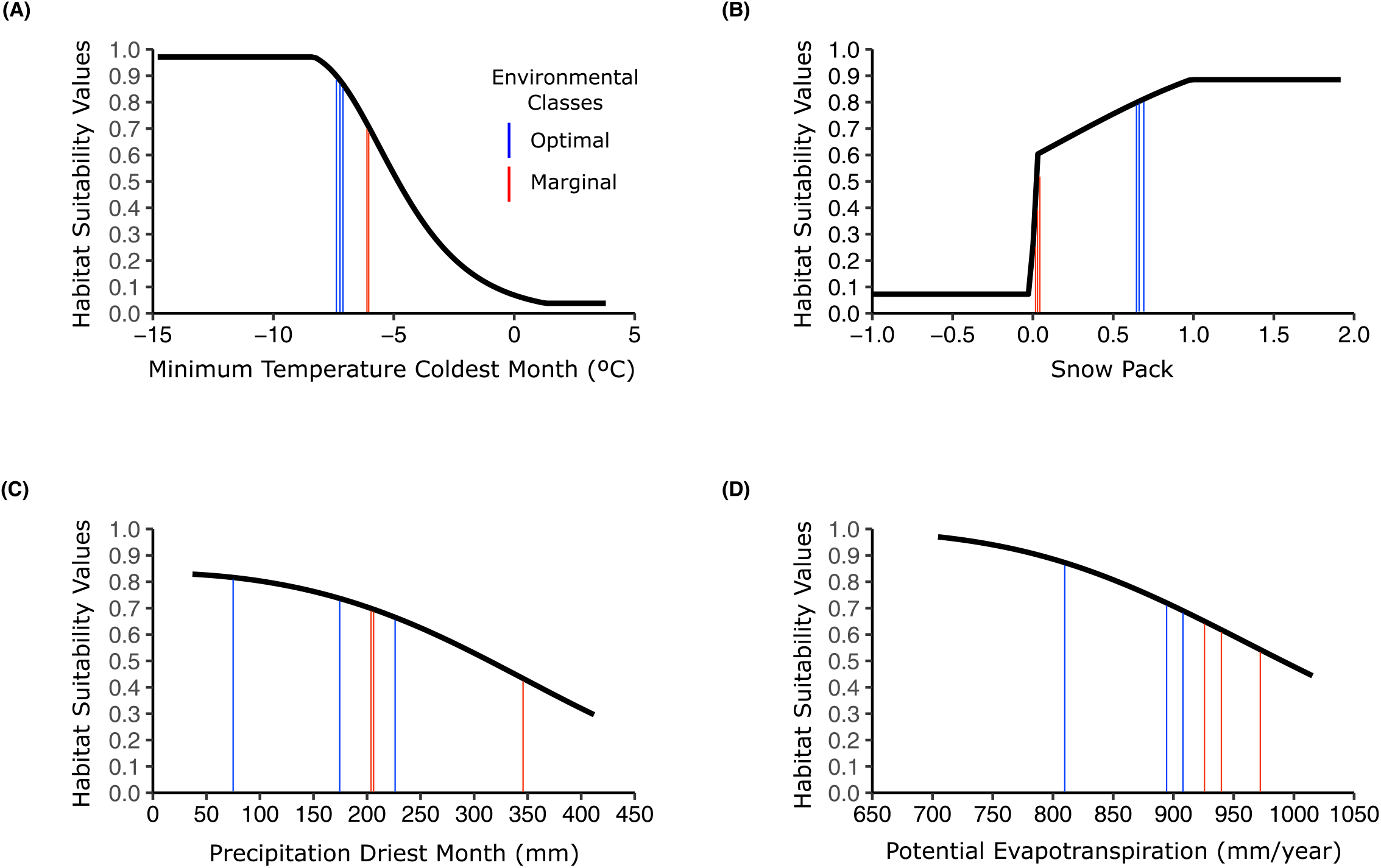
Response curves for the predictor variables used in MAXENT models of Mediterranean alpine species *Silene ciliata*. Y-axes show suitability values ranging from 0.0 (lowest) to 1.0 (highest). (A)Minimum temperature of the coldest month, (B) mean snow accumulation ratio in the thaw months, (C) precipitation and, (D) potential evapotranspiration in mm/year. Note that as minimum temperature of the coldest month increases, habitat suitability values decrease. Habitat suitability values are close to zero until snowpack values are positive (presences of snow cover). Precipitation of the driest month and potential evapotranspiration showed similar responses, with gradually decreasing HS values as both variables increased. Vertical bars represent values for each population for each variable. For detailed values see table A 5.1.

### 3.3 Functional traits

#### Categorical analysis

Environmental classification into optimal and marginal areas based on the ENM had a significant effect on all the phenological variables measured (Table 5). The covariate plant size only had a significant effect on flowering onset. Populations located in optimal areas displayed an earlier flowering onset, peak and end than populations inhabiting marginal areas (Table 5, Figure 3 A, B, C). Flowering season duration was narrower in populations from optimal areas (Figure 3 D, table S2). Onset showed a less differentiated response between optimal and marginal areas than flowering peak, duration and end (Table 5, Figure 3). In contrast, no significant associations were found between environmental classification and LT50_LOW_ or LT50_HIGH_ (Table 6).

**Table 5.** Analysis of the effect of environmental conditions and plant size on four phenological variables of *Silene ciliata* measured in a common garden experiment. In the quantitative approach, generalized linear models were used with habitat suitability values (HS) as fixed factors and plant size as a covariable. In the qualitative approach, Cox proportional hazards regression models were used with environmental classification (optimal, marginal) as fixed factors, population as random factor and plant size as a covariable.

| Continues approach |  |  |  |  |  | Categorical approach |  |  |  |  |  |
| --- | --- | --- | --- | --- | --- | --- | --- | --- | --- | --- | --- |
|  |  | Onset | Peak | End | Duration |  |  | Onset | Peak | End | Duration |
| | df | $\chi^2$ | $\chi^2$ | $\chi^2$ | $\chi^2$ | | df | $\chi^2$ | $\chi^2$ | $\chi^2$ | $\chi^2$ |
| Plant size | 1 | 6.4* | 2.4 | 2.0 | 5.9* | Plant size | 1 | 8.2** | 2.9 | 2.4 | 1.5 |
| HS values | 1 | 16.8*** | 27.3*** | 37.7*** | 16.6*** | Environmental<br>Classes | 1 | 3.9* | 19.7*** | 4.9* | 5.68* |
|  | R<sup>2</sup> | 0.32 | 0.40 | 0.42 | 0.38 |  | R<sup>2</sup> | 0.40 | 0.39 | 0.47 | 0.51 |
| *, P<0.05**, P<0.01; ***,P<0.001 |  |  |  |  |  |  |  |  |  |  |  |

\*, P<0.05\*\*, P<0.01; \*\*\*,P<0.001
HS values; habitat suitability values obtained from the ecological niche model, Environment; environmental classification in optimal and marginal areas based on HS values. Peak was defined as the number of days elapsed between the date of flowering onset of an individual and the date with the greatest number of open flowers on that individual.

**Table 6.**
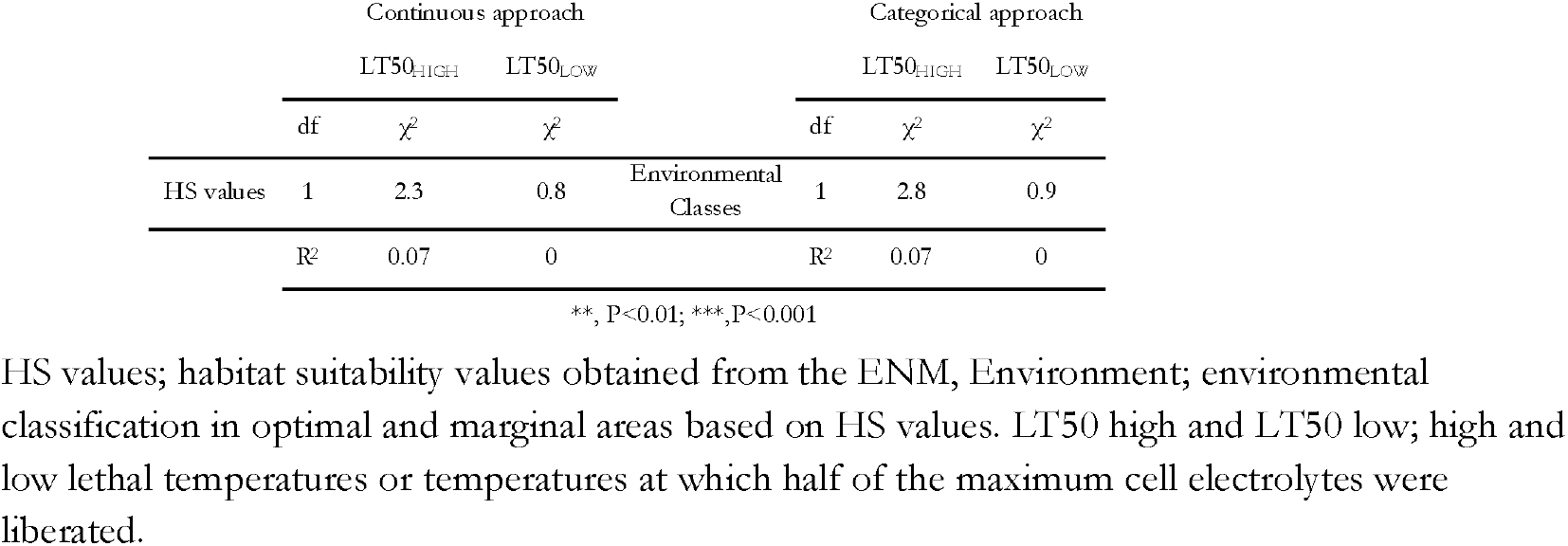
Analysis of the effect of environmental conditions on two cell temperature resistance variables (LT50_HIGH_ and LT50_LOW_) of *Silene ciliata*. In the quantitative approach, generalized linear models were used with habitat suitability values as fixed factors. In the qualitative approach, generalized linear models were used with environmental classification (optimal, marginal) as fixed factors and population as random factor.

**Figure 3.**
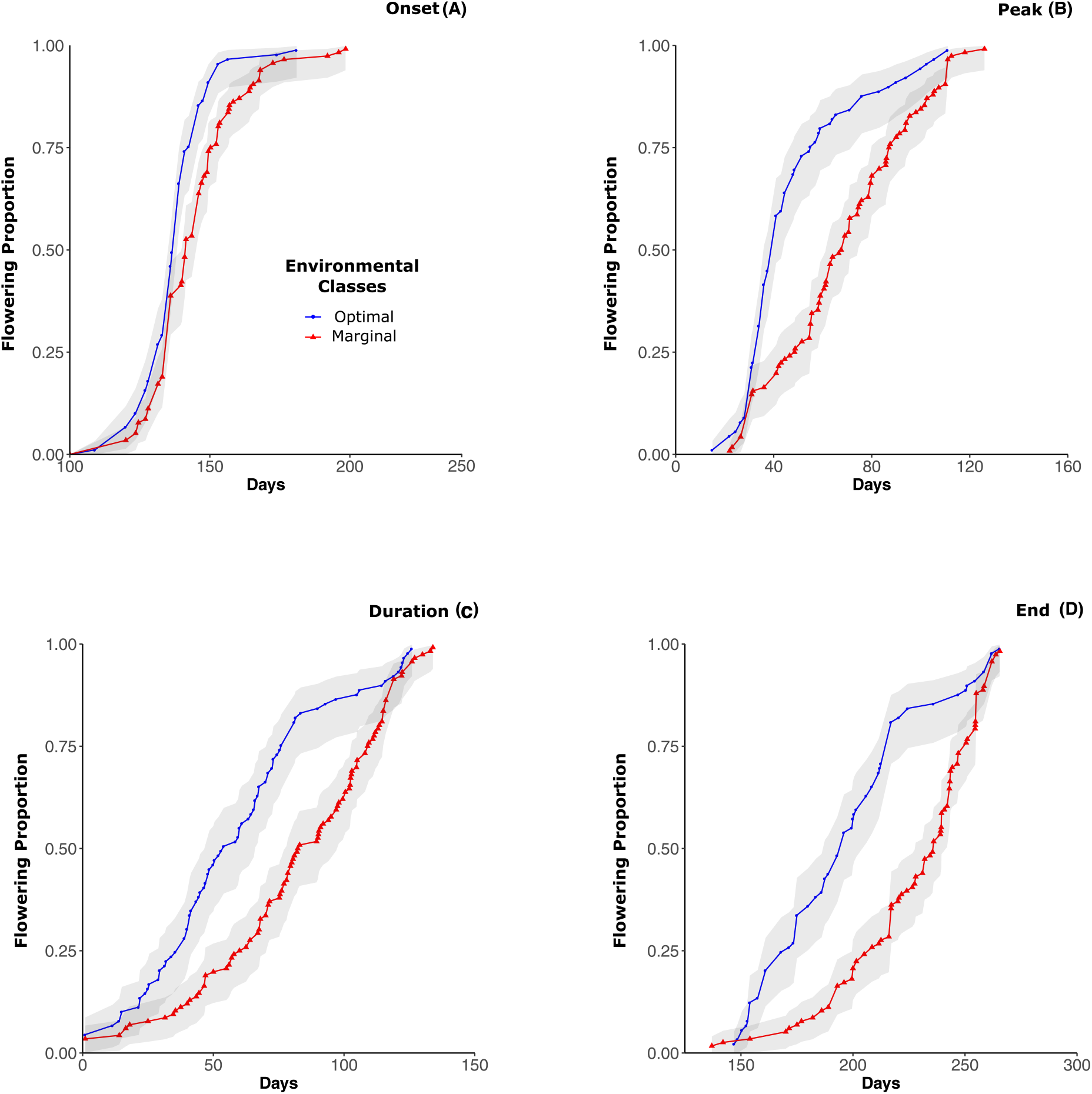
Effect of environmental classification (optimal and marginal areas) based on the habitat suitability values from the selected ENM on *Silene ciliata* phenological traits: Cumulative frequency curves of (A) flowering onset, (B) flowering peak, (C) flowering end and (D) flowering duration.

#### Continuous analysis

HS values from the ENM showed a significant negative relationship with each of the four flowering phenology variables (Table 5): as HS values increased, the values of onset, peak, duration and end decreased (Figure 4). Plant size had a significant positive effect on onset and a significant negative effect on duration. In contrast, as with the results for the categorical analyses, no significant associations were found between HS values and LT50_LOW_ or LT50_HIGH_ (Table 6).

**Figure 4.**
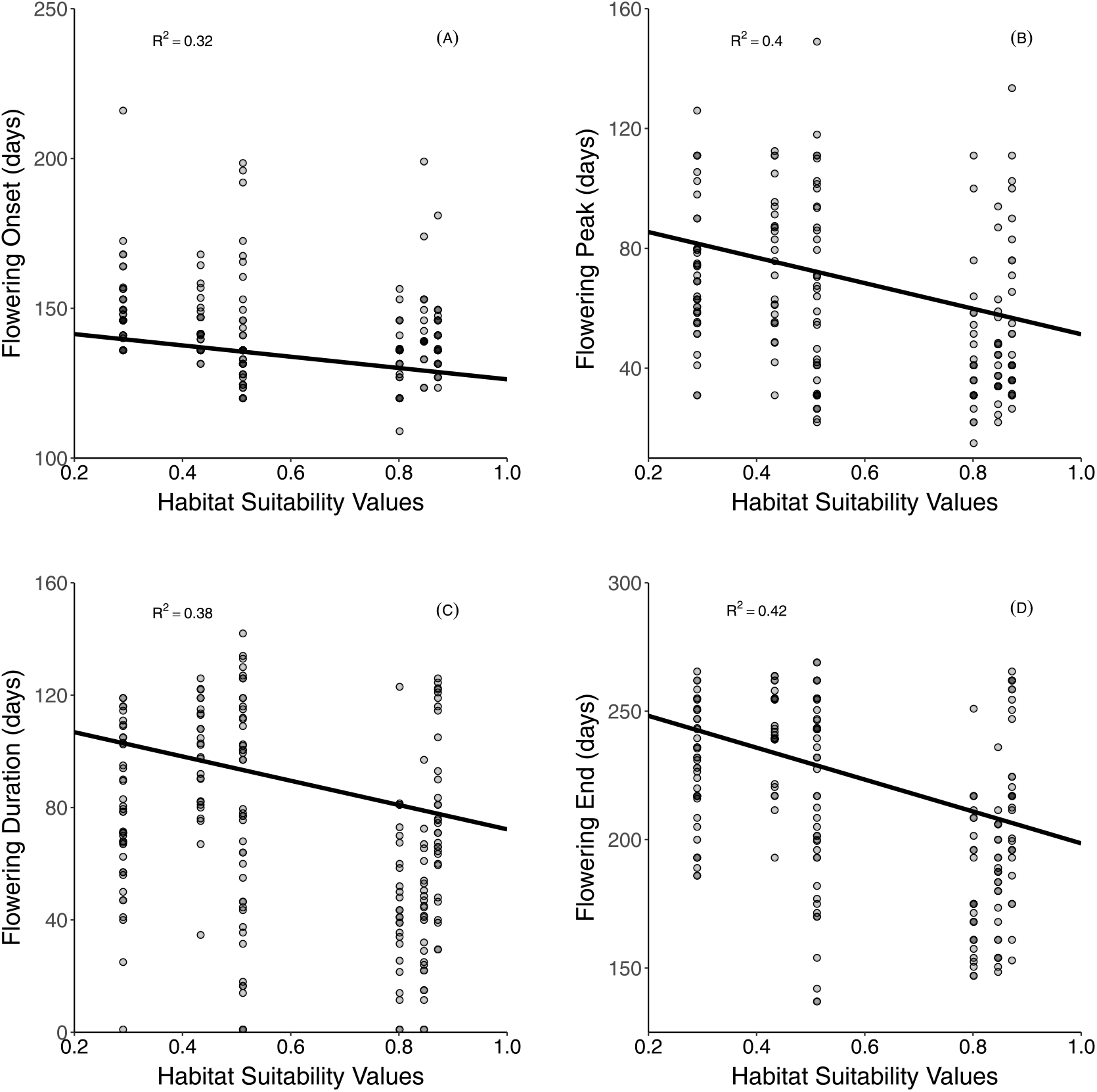
Relationship between habitat suitability values from the selected ENM and *Silene ciliata* phenological traits: (A) flowering onset, (B) flowering peak (B), (C) flowering end and (D) flowering duration.

## 4. Discussion

Results indicated functional genetic differentiation consistent with the predictions of environmental stressors within the *S. ciliata* distribution identified by the ENMs. We found genetically-based phenotypic differentiation in *phenological* traits between populations inhabiting areas classified as marginal vs. optimal based on ENMs and between populations with different HS values. In contrast, neither environmental classification into optimal and marginal areas nor raw HS values correlated with *physiological* traits, suggesting differential trait sensitivity to selective pressures imposed by differing environmental conditions.

Regarding the first point, the ENM was useful to distinguish different environmental conditions, to identify ecological marginality within the species range, and thus, it may also provide a valid approach to identify divergent selective pressures (Hällfors et al., 2016). Habitat suitability values generated by the model were closely associated to elevation, but also incorporated complex effects of aspect and topography. Indeed, strong elevation gradients in mountains create a complex environmental gradient in which different abiotic factors play different roles (Körner, 2003, 2007). The selected ENM was able to distinguish the relatively small-scale environmental variation experienced by *S. ciliata* (Figure 1). The main factors explaining species distribution and intra-range environmental differentiation were snowpack during thaw months and minimum temperature of coldest month. Snowpack variable combines information about temperature and precipitation at a particular time (López-Moreno et al., 2007), affecting timing and success of crucial life stages such as seed germination and plant flowering in *S. ciliata* (Giménez-Benavides et al., 2007a, 2011b) and other alpine plant species (Dunne et al., 2003; Giménez-Benavides et al., 2018; Inouye, 2008; Inouye et al., 2002; Lara-Romero et al., 2014a). Minimum temperature has been proposed as a reference variable to characterize environmental gradients and range limits in alpine ecosystems (Körner, 2003, 2007; Normand et al., 2009; Totland, 1999; Totland and Alatalo, 2002), reflecting alpine species requirement for low temperatures. Previous studies have documented differences in the needs of cold stratification between *S. ciliata* populations inhabiting different altitudes (García-Fernández et al., 2014; Giménez-Benavides et al., 2005), which may be related to the minimum temperature of the coldest month. Therefore, these two variables may be good proxies of the drivers that generate divergent selective pressures at the gradient extremes. Conversely, precipitation-related environmental variables generally show regional and variable patterns (Körner, 2003, 2007) and, as our model highlighted, these may be less representative of the environmental gradient in alpine areas.

The ENM also provided the basis to delimit environmental differentiation between optimal and marginal areas inside the species range. Previous studies with this and other alpine species have revealed marked ecological niche differences between populations inhabiting central and marginal areas inside their ranges related with fine-scale environmental variables (Giménez-Benavides et al., 2007b; Papuga et al., 2018). The ENM-based differentiation between optimal and marginal areas is congruent with demographic trends previously obtained (Giménez-Benavides et al., 2011a; Lara-Romero et al., 2014b, 2016a), where *S. ciliata* populations from optimal areas were at demographic equilibrium while those in marginal areas showed negative population growth rates. In this context ENMs are useful as hypothesis generators to study the ecological differentiation related with environmental marginality inside species distributions. Nevertheless, the extrapolation of these environmental classification to other areas of *S. ciliata* distribution or to other species should be taken with caution because ecological differentiation of plant populations related with abiotic factors is commonly species- and context-dependent (Leuschner et al., 2009; Papuga et al., 2018; Wagner et al., 2011).

Divergent natural selection is one of the main driving forces of genetic differentiation and thus adaptation (Kawecki, 2008; Kawecki and Ebert, 2004). Hence, the identification of genetically-based phenotypic differentiation between environmentally optimal and marginal areas provides relevant evidence in favor of the presence of divergent selective pressures. Both ENM-based optimal-marginal environmental classifications and HS values (categorical and continuous approaches) were able to detect genetic differentiation of phenological traits. Populations from optimal areas (higher HS values) flowered earlier and faster than populations inhabiting marginal areas (lower HS values) when grown in common garden settings. This is congruent with previous results with the co-occurring plant *Armeria caespitosa* in the same ecosystems (Lara-Romero et al., 2014a). The genetically-based phenological differentiation we found could also result from neutral processes separate from selection (Garcia-Ramos and Kirkpatrick, 1997). Nevertheless, phenological reproductive traits are likely to be under strong selection since they have been shown to be critical for plant fitness (Fox, 2003; Levin, 2006), capable of rapidly evolving under climatic variation (Franks et al., 2007) and influencing species distributions and potential responses to climate change (Banta et al., 2012; Chuine, 2010). This is in agreement with empirical studies showing that these traits generally show high evolvability (Kawakami et al., 2011; Méndez-Vigo et al., 2013; Volis, 2011) and with a meta-analysis showing consistent selection patterns (Munguía-Rosas et al., 2011). Moreover, selection analyses performed under field conditions by Giménez-Benavides et al. (2011b) for the same species in the same areas showed that flowering onset and duration are under selection in populations at different elevations (i.e. selected for early flowering) (see also chapter 3). In this vein, a parallel transcriptomic study using material collected from the same study’s populations identified a set of candidate SNPs with unusually high allele frequency differentiation between marginal and optimal environments (Sacristán-Bajo et al., 2019). Many of these SNPs derived from candidate genes in the flower development pathway (Sacristán-Bajo et al., 2019). Thus, these selection analysis and genomic approaches provide increased evidence that genetically-based phenological differentiation found in this study is the result of divergent selection between marginal and optimal populations.

In our study no differentiation of the tested physiological traits was found, which is in line with previous research with this species and others in the same ecosystem (Pescador et al., 2015).Thus, our results support the idea that phenological traits are more sensitive to divergent selection than physiological traits (Bradshaw and Holzapfel, 2008; Hoffmann and Sgró, 2011). A possible explanation to this is that changing the phenology of an organism in response to an environmental pressure is an escape mechanism that is easier to carry out than changing the physiological response needed to adapt to the new environmental conditions (Bradshaw and Holzapfel, 2008, 2010; Hoffmann and Sgró, 2011; Kovach et al., 2012).

Our findings, resulting from the joint use of ENMs and functional traits, experimentally supported the hypothesis that marginality—expressed by habitat suitability differentiation—is an important factor predicting variation of functional traits among populations (Lira-Noriega and Manthey, 2014; Madeira de Madeiros et al., 2018). In this context, the ecological uniqueness of environmentally marginal populations highlights their evolutionary potential and conservation value (Papuga et al., 2018). Considering that the development of evolutionary adaptations to a marginal habitat may require substantial levels of within-population genetic diversity, it must be noted that no clear relationship between genetic diversity loss and marginality has been found, showing context-dependent performances (Diniz-Filho et al., 2009; Duncan et al., 2015; Lira-Noriega and Manthey, 2014). Thus, species history plays a key role in changes to range limits and thus in genetic diversity variation in the CPH framework (Duncan et al., 2015; Hampe and Petit, 2005).

The case of *S. ciliata* highlights the value of conserving marginal populations (Channell, 2004) based on their genetically-based phenotypic differentiation. Nevertheless, further research is needed to confirm adaptive processes occurring in marginal populations and the adaptive value of such phenotypic variation: This could include (i) the evaluation of genetic diversity and structure possibly related with environmental divergence, (ii) evaluation of traits under selection, (iii) direct local adaptation evaluation using field or genomics approaches (e.g. reciprocal transplants, gene expression) and (iv) the study of gene flow effects on population performance, among others.

### Conclusions

The ENM developed was useful to characterize the environmental niche experienced by the different studied populations of *S. ciliata*, to define environmental marginality, and potential environment-induced selective pressures. The use of ENMs as hypothesis generators regarding selective pressures, subsequently tested with phenotypic traits evaluated under common garden conditions, can be a powerful tool to disentangle divergent selection inside species distributions. If our final aim is to understand the best possible physiological and evolutionary capabilities of species under a context of climate change, a modeling approach in conjunction with laboratory and field experiments is recommended (Dixon and Busch, 2017; Swab et al., 2015). In this context, after a previous use of correlative models as hypothesis generators and testing via experimental studies such as this one, mechanistic models may be a powerful approach to account for intra-range species phenotypic variability and improve climate change species response studies (Benito-Garzón et al., 2011, 2019; Valladares et al., 2014). Since the adaptive potential of marginal populations is most likely species- and context-dependent, our approach provides a simple and practical way to study adaptive processes for each species and circumstance and to provide insights about the adaptive potential and conservation value of their marginal populations. Hopefully, such a line of research will lead to general conclusions regarding different classes of organisms and ecosystems. Further research should be undertaken to ensure the adaptive potential of *S. ciliata* populations inhabiting marginal areas.

## Supporting information

Supplementary material including tables, figures caption and more detailed information related to the main text

## References

Abeli, T., Gentili, R., Mondoni, A., Orsenigo, S., and Rossi, G. (2014). Effects of marginality on plant population performance. J. Biogeogr. 41, 239–249. doi:10.1111/jbi.12215.

Abeli, T., and Orsenigo, S. (2018). The importance of marginal population hotspots of cold-adapted species for research on climate change and conservation. J. Biogeogr. 45, 977–985. doi:10.1111/jbi.13196.

Aiello□Lammens, M. E., Boria, R. A., Radosavljevic, A., Vilela, B., and Anderson, R. P. (2015). spThin: an R package for spatial thinning of species occurrence records for use in ecological niche models. Ecography (Cop.). 38, 541–545.

Anderson, R. P. (2012). Harnessing the world’s biodiversity data□: promise and peril in ecological niche modeling of species distributions. Ann. N. Y. Acad. Sci., 1–15. doi:10.1111/j.1749-6632.2011.06440.x.

Anderson, R. P. (2013). A framework for using niche models to estimate impacts of climate change on species distributions. Ann. N. Y. Acad. Sci. 1297, 8–28. doi:10.1111/nyas.12264.

Anderson, R. P., and Gonzalez, I. (2011). Species-specific tuning increases robustness to sampling bias in models of species distributions: An implementation with Maxent. Ecol. Modell. 222, 2796–2811. doi:10.1016/j.ecolmodel.2011.04.011.

Anderson, R. P., and Raza, A. (2010). The effect of the extent of the study region on GIS models of species geographic distributions and estimates of niche evolution: Preliminary tests with montane rodents (genus Nephelomys) in Venezuela. J. Biogeogr. 37, 1378–1393. doi:10.1111/j.1365-2699.2010.02290.x.

Angert, A. L. (2006). Demographyof central and marginal populations of monkeyflowers (Mimulus cardinalis and M. lewisii). Ecology 87, 2014–2025.

Araújo, M. B., Anderson, R. P., Barbosa, A. M., and Beale, C. M. (2019). Standards for distribution models in biodiversity assessments. Sci. Adv. 5, 1–10.

Araujo, M. B., and Guisan, A. (2006). Five (or so) challenges for species distribution modelling. J. Biogeogr. 33, 1677–1688. doi:10.1111/j.1365-2699.2006.01584.x.

Banta, J. A., Ehrenreich, I. M., Gerard, S., Chou, L., Wilczek, A., Schmitt, J., et al. (2012). Climate envelope modelling reveals intraspecific relationships among flowering phenology, niche breadth and potential range size in Arabidopsis thaliana. Ecol. Lett. 15, 769–777. doi:10.1111/j.1461-0248.2012.01796.x.

Baskin, C. C., and Baskin, J. M. (1998). Seeds: ecology, biogeography, and, evolution of dormancy and germination. San Diego, USA: Elsevier.

Bates, D., Mächler, M., Bolker, B., and Walker, S. (2014). Fitting linear mixed-effects models using lme4. arXiv Prepr. arXiv1406.5823.

Benito-Garzón, M., Alía, R., Robson, T. M., and Zavala, M. A. (2011). Intra-specific variability and plasticity influence potential tree species distributions under climate change. Glob. Ecol. Biogeogr. 20, 766–778. doi:10.1111/j.1466-8238.2010.00646.x.

Benito-Garzón, M., Robson, T. M., and Hampe, A. (2019). ATraitSDMs□: species distribution models that account for local adaptation and phenotypic plasticity. New Phytol., 1–9. doi:10.1111/nph.15716.

Blanquart, F., Kaltz, O., and Gandon, S. (2013). A practical guide to measuring local adaptation. Ecol. Lett. 16, 1195–1205. doi:10.1111/ele.12150.

Boria, R. A., Olson, L. E., Goodman, S. M., and Anderson, R. P. (2014). Spatial filtering to reduce sampling bias can improve the performance of ecological niche models. Ecol. Modell. 275, 73–77. doi:10.1016/j.ecolmodel.2013.12.012.

Bradshaw, W. E., and Holzapfel, C. M. (2008). Genetic response to rapid climate change□: it’s seasonal timing that matters. Mol. Ecol. 17, 157–166. doi:10.1111/j.1365-294X.2007.03509.x.

Bradshaw, W. E., and Holzapfel, C. M. (2010). Light, Time, and the Physiology of Biotic Response to Rapid Climate Change in Animals. Annu. Rev. Physiol. 72, 147–166. doi:10.1146/annurev-physiol-021909-135837.

Brown, J. H. (1984). On the Relationship between Abundance and Distribution of Species. Am. Nat. 124, 255–279. doi:10.1086/284267.

Brussard, P. F. (1984). Geographic patterns and environmental gradients: the central-marginal model in Drosophila revisited. Annu. Rev. Ecol. Syst. 15, 25–64.

Channell, R. (2004). The conservation value of peripheral populations: the supporting science. in Proceedings of the species at risk 2004 pathways to recovery conference (Species at Risk 2004 Pathways to Recovery Conference Organizing Committee …), 1–17.

Chuine, I. (2010). Why does phenology drive species distribution? Philos. Trans. R. Soc. B Biol. Sci. 365, 3149–3160. doi:10.1098/rstb.2010.0142.

Didden-Zopfy, B., and Nobel, P. S. (1982). High Temperature Tolerance and Heat Acclimation of Opuntia bigelovii. Oecologia 52, 176–180.

Diniz-Filho, J. A. F., Nabout, J. C., Bini, L. M., Soares, T. N., de Campus Telles, M. P., de Marco, P., et al. (2009). Niche modelling and landscape genetics of Caryocar brasiliense (“Pequi” tree: Caryocaraceae) in Brazilian Cerrado: An integrative approach for evaluating central-peripheral population patterns. Tree Genet. Genomes 5, 617–627. doi:10.1007/s11295-009-0214-0.

Dixon, A. L., and Busch, W. (2017). Common garden test of range limits as predicted by a species distribution model in the annual plant Mimulus. Am. J. Bot. 104, 817–827. doi:10.3732/ajb.1600414.

Drennan, P. M. (2009). “Temperature influences on plant species of arid a semi-arid regions with emphasis on CAM succulents,” in Perspectives in Biophysical Plant Ecophysiology, eds. E. Barrera and W. K. Smith (Mexico City, Mexico: UNAM), 57–94.

Duncan, S. I., Crespi, E. J., Mattheus, N. M., and Rissler, L. J. (2015). History matters more when explaining genetic diversity within the context of the core-periphery hypothesis. Mol. Ecol. 24, 4323–4336. doi:10.1111/mec.13315.

Dunne, J. A., Harte, J., and Taylor, K. J. (2003). Subalpine Meadow Flowering Phenology Responses to Climate Change□: Integrating Experimental and Gradient Methods Author (s): Jennifer A. Dunne, John Harte and Kevin J. Taylor Published by□: Wiley on behalf of the Ecological Society of America Stable. Ecol. Monogr. 73, 69–86.

Eckert, C. G., Samis, K. E., and Lougheed, S. C. (2008). Genetic variation across species’ geographical ranges: The central-marginal hypothesis and beyond. Mol. Ecol. 17, 1170–1188. doi:10.1111/j.1365-294X.2007.03659.x.

Elith, J., and Leathwick, J. R. (2009). Species Distribution Models□: Ecological Explanation and Prediction Across Space and Time. Annu. Rev. Ecol. Evol. Syst. 40, 677–697. doi:10.1146/annurev.ecolsys.110308.120159.

Escudero, A., Giménez-Benavides, L., Iriondo, J. M., and Rubio, A. (2004). Patch dynamics and islands of fertility in a high mountain Mediterranean community. Arctic, Antarct. Alp. Res. 36, 518–527. doi:10.1657/1523-0430(2004)036[0518:PDAIOF]2.0.CO;2.

Fox, G. A. (2003). Assortative mating and plant phenology: Evolutionary and practical consequences. Evol. Ecol. Res. 5, 1–18.

Fox, J., and Weisberg, S. (2011). An R companion to applied regression. Sage Publications.

Franks, S. J., Sim, S., and Weis, A. E. (2007). Rapid evolution of flowering time by an annual plant in response to a climate fluctuation. PNAS 104, 1278–1282.

Galante, P. J., Alade, B., Muscarella, R., Jansa, S. A., Goodman, S. M., and Anderson, R. P. (2017). The challenge of modeling niches and distributions for data-poor species□: a comprehensive approach to model complexity. Ecography (Cop.). 40, 1–10. doi:10.1111/ecog.02909.

García-Fernández, A., Escudero, A., Lara-Romero, C., and Iriondo, J. M. (2014). Effects of the duration of cold stratification on early life stages of the Mediterranean alpine plant Silene ciliata. Plant Biol. 17, 344–350. doi:10.1111/plb.12226.

García-Fernández, A., Iriondo, J. M., Bartels, D., and Garcı, A. (2013). Response to artificial drying until drought-induced death in different elevation populations of a high-mountain plant. Plant Biol. 15, 93–100. doi:10.1111/j.1438-8677.2012.00638.x.

García-Fernández, A., Segarra-Moragues, J. G., Widmer, A., Escudero, A., and Iriondo, J. M. (2012). Unravelling genetics at the top: Mountain islands or isolated belts? Ann. Bot. 110, 1221–1232. doi:10.1093/aob/mcs195.

Garcia-Ramos, G., and Kirkpatrick, M. (1997). Genetic Models of Adaptation and Gene Flow in Peripheral Populations. Evolution (N. Y). 51, 21. doi:10.2307/2410956.

Gaston, K. J., and Sheffield, S. (2009). Geographic range limits□: achieving synthesis. Proc. R. Soc. B Biol. Sci. 276, 1395–1406. doi:10.1098/rspb.2008.1480.

Giménez-Benavides, L., Albert, M. J., Iriondo, J. M., and Escudero, A. (2011a). Demographic processes of upward range contraction in a long-lived Mediterranean high mountain plant. Ecography (Cop.). 34, 85–93. doi:10.1111/j.1600-0587.2010.06250.x.

Giménez-Benavides, L., Escudero, A., García-Camacho, R., García-Fernández, A., Iriondo, J. M., Lara-Romero, C., et al. (2018). How does climate change affect regeneration of Mediterranean high-mountain plants? An integration and synthesis of current knowledge. Plant Biol. 20, 50–62. doi:10.1111/plb.12643.

Giménez-Benavides, L., Escudero, A., and Iriondo, J. M. (2007a). Local adaptation enhances seedling recruitment along an altitudinal gradient in a high mountain mediterranean plant. Ann. Bot. 99, 723–734. doi:10.1093/aob/mcm007.

Giménez-Benavides, L., Escudero, A., and Iriondo, J. M. (2007b). Reproductive limits of a late-flowering high-mountain Mediterranean plant along an elevational climate gradient. New Phytol. 173, 367–382. doi:10.1111/j.1469-8137.2006.01932.x.

Giménez-Benavides, L., Escudero, A., and Pérez-García, F. (2005). Seed germination of high mountain Mediterranean species: Altitudinal, interpopulation and interannual variability. Ecol. Res. 20, 433–444. doi:10.1007/s11284-005-0059-4.

Giménez-Benavides, L., García-Camacho, R., Iriondo, J. M., and Escudero, A. (2011b). Selection on flowering time in Mediterranean high-mountain plants under global warming. Evol. Ecol. 25, 777–794. doi:10.1007/s10682-010-9440-z.

Grinnell, J. (1917). The niche-relationships of the California Thrasher. Auk 34, 427–433.

Guevara, L., Gerstner, B. E., Kass, J. M., and Anderson, R. P. (2018). Toward ecologically realistic predictions of species distributions: A cross-time example from tropical montane cloud forests. Glob. Chang. Biol. 24, 1511–1522. doi:10.1111/gcb.13992.

Guisan, A., and Thuiller, W. (2005). Predicting species distribution□: offering more than simple habitat models. 8, 993–1009. doi:10.1111/j.1461-0248.2005.00792.x.

Hällfors, M. H., Liao, J., Dzurisin, J., Grundel, R., Hyvärinen, M., Towle, K., et al. (2016). Addressing potential local adaptation in species distribution models: Implications for conservation under climate change. Ecol. Appl. 26, 1154–1169. doi:10.1890/15-0926.

Hampe, A., and Petit, R. J. (2005). Conserving biodiversity under climate change: The rear edge matters. Ecol. Lett. 8, 461–467. doi:10.1111/j.1461-0248.2005.00739.x.

Hardie, D. C., and Hutchings, J. A. (2010). Evolutionary ecology at the extremes of species’ ranges. Environ. Rev. 18, 1–20. doi:10.1139/A09-014.

Hengeveld, R., and Haeck, J. (1982). The distribution of abundance. I. Measurements. J. Biogeogr. 9, 303–316.

Herlihy, C. R., and Eckert, C. G. (2005). Evolution of self-fertilization at geographic range margins? A comparison of demographic, floral, and mating system variables in central vs. peripheral populations of Aquilegia Canadensis (Ranunculaceae). Am. J. Bot. 92, 744–751.

Herrera, C. M., and Bazaga, P. (2008). Population-genomic approach reveals adaptive floral divergence in discrete populations of a hawk moth-pollinated violet. Mol. Ecol. 17, 5378–5390. doi:10.1111/j.1365-294X.2008.04004.x.

Hoffmann, A. A., and Blows, M. W. (1994). Reviews 12. Trends Ecol. Evol. 9, 223–227.

Hoffmann, A. A., and Sgró, C. M. (2011). Climate change and evolutionary adaptation. Nature 470, 479–485. doi:10.1038/nature09670.

Holt, R. D., and Keitt, T. H. (2005). Species’ Borders□: A Unifying Theme in Ecology. Oikos 108, 3–6.

Hunter, M. L., and Hutchinson, A. (1994). The Virtues and Shortcomings of Parochialism□: Conserving Species That Are Locally Rare, but Globally Common. Conserv. Biol. 8, 1163–1165.

Hutchinson, G. E. (1957). Concluding remarks Cold Spring Harbor Symposia on Quantitative Biology. GS SEARCH 22, 415–427.

Inouye, D. W. (2008). Effects of Climate Change on Phenology, Frost Damage and Floral Abundance of Montane Wild Flowers. America (NY). 89, 353–362.

Inouye, D. W., Morales, M. A., and Dodge, G. J. (2002). Variation in timing and abundance of flowering by Delphinium barbeyi Huth (Ranunculaceae): the roles of snowpack, frost, and La Niña, in the context of climate change. 543–550. doi:10.1007/s00442-001-0835-y.

Kawakami, T., Morgan, T. J., Nippert, J. B., Ocheltree, T. W., Keith, R., Dhakal, P., et al. (2011). Natural selection drives clinal life history patterns in the perennial sunflower species, Helianthus maximiliani. Mol. Ecol. 20, 2318–2328. doi:10.1111/j.1365-294X.2011.05105.x.

Kawecki, T. J. (2008). Adaptation to Marginal Habitats. Annu. Rev. Ecol. Evol. Syst. 39, 321–342. doi:10.1146/annurev.ecolsys.38.091206.095622.

Kawecki, T. J., and Ebert, D. (2004). Conceptual issues in local adaptation. Ecol. Lett. 7, 1225–1241. doi:10.1111/j.1461-0248.2004.00684.x.

Kearney, M., and Porter, W. (2009). Mechanistic niche modelling: Combining physiological and spatial data to predict species’ ranges. Ecol. Lett. 12, 334–350. doi:10.1111/j.1461-0248.2008.01277.x.

Körner, C. (2003). Alpine Plant Life. Functional Plant Ecology of High Mountain Ecosystems. Berlin: Springer-Verlag Berlin Heidelberg doi:10.1007/978-3-642-18970-8.

Körner, C. (2007). The use of “altitude” in ecological research. Trends Ecol. Evol. 22, 569–574. doi:10.1016/j.tree.2007.09.006.

Kovach, R. P., Gharrett, A. J., and Tallmon, D. A. (2012). Genetic change for earlier migration timing in a pink salmon population. Proc. R. Soc. B Biol. Sci. 279, 3870–3878. doi:10.1098/rspb.2012.1158.

Kyrkou, I., Iriondo, J. M., and García-Fernández, A. (2015). A glacial survivor of the alpine Mediterranean region: phylogenetic and phylogeographic insights into Silene ciliata Pourr. (Caryophyllaceae). PeerJ 3, 1–19. doi:10.7717/peerj.1193.

Lara-Romero, C., de la Cruz, M., Escribano-Ávila, G., García-Fernández, A., and Iriondo, J. M. (2016a). What causes conspecific plant aggregation? Disentangling the role of dispersal, habitat heterogeneity and plant–plant interactions. Oikos 125, 1304–1313. doi:10.1111/oik.03099.

Lara-Romero, C., García-Camacho, R., Escudero, A., and Iriondo, J. M. (2014a). Genetic variation in flowering phenology and reproductive performance in a Mediterranean high-mountain specialist, Armeria caespitosa (Plumbaginaceae). Bot. J. Linn. Soc. 176, 384–395. doi:10.1111/boj.12208.

Lara-Romero, C., García-Fernández, A., Robledo-Arnuncio, J. J., Roumet, M., Morente-López, J., López-Gil, A., et al. (2016b). Individual spatial aggregation correlates with between-population variation in fine-scale genetic structure of Silene ciliata (Caryophyllaceae). Heredity (Edinb). 116. doi:10.1038/hdy.2015.102.

Lara-Romero, C., García-Fernández, A., Robledo-Arnuncio, J. J., Roumet, M., Morente-López, J., López-Gil, A., et al. (2016c). Individual spatial aggregation correlates with between-population variation in fine-scale genetic structure of Silene ciliata (Caryophyllaceae). Heredity (Edinb). 116, 417–423. doi:10.1038/hdy.2015.102.

Lara-Romero, C., Robledo-Arnuncio, J. J., García-Fernández, A., and Iriondo, J. M. (2014b). Assessing intraspecific variation in effective dispersal along an altitudinal gradient: A test in two Mediterranean high-mountain plants. PLoS One 9. doi:10.1371/journal.pone.0087189.

Lesica, P., and Allendorf, F. W. (1995). When Are Peripheral Populations Valuable for Conservation□? Conserv. Biol. 9, 753–760.

Leuschner, C., Köchemann, B., and Buschmann, H. (2009). Forest Ecology and Management Abundance, niche breadth, and niche occupation of Central European tree species in the centre and at the margin of their distribution range. For. Ecol. 258, 1248–1259. doi:10.1016/j.foreco.2009.06.020.

Levin, D. A. (2006). Flowering Phenology in Relation to Adaptive Radiation. Syst. Bot. 31, 239–246. doi:10.1600/036364406777585928.

Lira-Noriega, A., and Manthey, J. D. (2014). Relationship of genetic diversity and niche centrality: A survey and analysis. Evolution (N. Y). 68, 1082–1093. doi:10.1111/evo.12343.

López-Moreno, J. I., Vicente-Serrano, S. M., and Lanjeri, S. (2007). Mapping snowpack distribution over large areas using GIS and interpolation techniques. Clim. Res. 33, 257.

Luceño, M., Vargas, P., and others (1991). Guía Botánica del Sistema Central español.

Madeira de Madeiros, C., Hernández-Lambraño, R. E., Felix Ribeiro, K. A., and Sánchez Agudo, J. Á. (2018). Living on the edge□: do central and marginal populations of plants differ in habitat suitability□? Plant Ecol. 219, 1029–1043.

Méndez-Vigo, B., Gomaa, N. H., Alonso-Blanco, C., and Xavier Picó, F. (2013). Among- and within-population variation in flowering time of Iberian Arabidopsis thaliana estimated in field and glasshouse conditions. New Phytol. 197, 1332–1343. doi:10.1111/nph.12082.

Merow, C., Smith, M. J., Edwards, T. C., Guisan, A., Mcmahon, S. M., Normand, S., et al. (2014). What do we gain from simplicity versus complexity in species distribution models? Ecography (Cop.). 37, 1267–1281. doi:10.1111/ecog.00845.

Millar, C. I., and Libby, W. J. (1991). Strategies for conserving clinal, ecotypic, and disjunct population diversity in widespread species. Genet. Conserv. rare plants 149, 170.

Moritz, C., and Agudo, R. (2013). The Future of Species Under Climate. Science 341, 504–507. doi:10.1126/science.1237190.

Munguía-Rosas, M. A., Ollerton, J., Parra-Tabla, V., and De-Nova, J. A. (2011). Meta-analysis of phenotypic selection on flowering phenology suggests that early flowering plants are favoured. Ecol. Lett. 14, 511–521. doi:10.1111/j.1461-0248.2011.01601.x.

Muscarella, R., Galante, P. J., Soley□Guardia, M., Boria, R. A., Kass, J. M., Uriarte, M., et al. (2014). ENM eval: An R package for conducting spatially independent evaluations and estimating optimal model complexity for Maxent ecological niche models. Methods Ecol. Evol. 5, 1198–1205.

Ninyerola, M., Pons, X., and Roure, J. M. (2005). Atlas climático digital de la Península Ibérica: metodología y aplicaciones en bioclimatología y geobotánica. Universitat Autònoma de Barcelona, Departament de Biologia Animal, Biologia Vegetal i Ecologia (Unitat de Botánica).

Nobel, P. S., Barrera, E. De, Beilman, D. W., Doherty, J. H., Zutta, R., Nobel, P. S., et al. (2002). Temperature Limitations for Cultivation of Edible Cacti in California. Calif. Bot. Soc. 49, 228–236.

Normand, S., Treier, U. A., Randin, C., Vittoz, P., Guisan, A., and Svenning, J. C. (2009). Importance of abiotic stress as a range-limit determinant for European plants: Insights from species responses to climatic gradients. Glob. Ecol. Biogeogr. 18, 437–449. doi:10.1111/j.1466-8238.2009.00451.x.

Owens, H. L., Campbell, L. P., Dornak, L. L., Saupe, E. E., Barve, N., Soberón, J., et al. (2013). Constraints on interpretation of ecological niche models by limited environmental ranges on calibration areas. Ecol. Modell. 263, 10–18. doi:10.1016/j.ecolmodel.2013.04.011.

Papuga, G., Gauthier, P., Pons, V., Farris, E., and Thompson, J. D. (2018). Ecological niche differentiation in peripheral populations: A comparative analysis of eleven Mediterranean plant species. Ecography (Cop.)., 1–15. doi:10.1111/ecog.03331.

Pescador, D. S., De Bello, F., Valladares, F., and Escudero, A. (2015). Plant trait variation along an altitudinal gradient in mediterranean high mountain grasslands: Controlling the species turnover effect. PLoS One 10, 1–16. doi:10.1371/journal.pone.0118876.

Peterson, A. T., Soberón, J., Pearson, R. G., Anderson, R. P., Martínez-Meyer, E., Nakamura, M., et al. (2011). Ecological niches and geographic distributions Princeton University Press. Princeton, NJ 328pp.

Phillips, S. J. (2011). “A brief tutorial on Maxent,” in AT&T Research (Princeton,).

Phillips, S. J., Anderson, R. P., and Schapire, R. E. (2006). Maximum entropy modeling of species geographic distributions. Ecol. Modell. 190, 231–259. doi:10.1016/j.ecolmodel.2005.03.026.

Phillips, S. J., and Dudík, M. (2008). Modeling of species distributions with Maxent□: new extensions and a comprehensive evaluation. Ecography (Cop.). 31, 161–175. doi:10.1111/j.2007.0906-7590.05203.x.

Pironon, S., Papuga, G., Villellas, J., Angert, A. L., García, M. B., and Thompson, J. D. (2017). Geographic variation in genetic and demographic performance: new insights from an old biogeographical paradigm. Biol. Rev. 92, 1877–1909. doi:10.1111/brv.12313.

Pironon, S., Villellas, J., Morris, W. F., Doak, D. F., and García, M. B. (2015). Do geographic, climatic or historical ranges differentiate the performance of central versus peripheral populations? Glob. Ecol. Biogeogr. 24, 611–620. doi:10.1111/geb.12263.

Pouget, M., Youssef, S., Migliore, J., Juin, M., Médail, F., and Baumel, A. (2013). Phylogeography sheds light on the central – marginal hypothesis in a Mediterranean narrow endemic plant. Ann. Bot. 112, 1409–1420. doi:10.1093/aob/mct183.

Radosavljevic, A., and Anderson, R. P. (2014). Making better Maxent models of species distributions: Complexity, overfitting and evaluation. J. Biogeogr. 41, 629–643. doi:10.1111/jbi.12227.

Rivas-Martínez, S., Fernández-González, F., Sánchez-Mata, D., and Pizarro, J. M. (1990). Vegetación de la Sierra de Guadarrama. Itinera Geobot. 4, 3–132.

Roberts, D. R., Bahn, V., Ciuti, S., Boyce, M. S., Elith, J., Guillera-arroita, G., et al. (2017). Cross-validation strategies for data with temporal, spatial, hierarchical, or phylogenetic structure. Ecography (Cop.). 40, 913–929. doi:10.1111/ecog.02881.

Rolland, J., Lavergne, S., and Manel, S. (2015). Combining niche modelling and landscape genetics to study local adaptation: A novel approach illustrated using alpine plants. Perspect. Plant Ecol. Evol. Syst. 17, 491–499. doi:10.1016/j.ppees.2015.07.005.

Sacristán-Bajo, S., García-Fernández, A., Iriondo, J. M., and Lara-Romero, C. (2019). Transcriptome assembly and polymorphism detection in Silene ciliata (Caryophyllaceae). Plant Genet. Resour., 1–4.

Saupe, E. E., Barve, V., Myers, C. E., Soberón, J., Barve, N., Hensz, C. M., et al. (2012). Variation in niche and distribution model performance□: The need for a priori assessment of key causal factors. Ecol. Modell. 237–238, 11–22. doi:10.1016/j.ecolmodel.2012.04.001.

Searcy, C. A., and Shaffer, H. B. (2016). Do Ecological Niche Models Accurately Identify Climatic Determinants of Species Ranges□? Am. Nat. 187, 423–435. doi:10.1086/685387.

Sexton, J. P., McIntyre, P. J., Angert, A. L., and Rice, K. J. (2009). Evolution and ecology of species range limits. Annu. Rev. Ecol. Evol. Syst. 40, 415–436. doi:10.1146/annurev.ecolsys.110308.120317.

Soto-Correa, J. C., Saenz-Romero, C., Lindig-Cisneros, R., and de la Barrera, E. (2013). The neotropical shrub Lupinus elegans, from temperate forests, may not adapt to climate change. Plant Biol. 15, 607–610. doi:10.1111/j.1438-8677.2012.00716.x.

Soule, M. (1973). The epistasis cycle: a theory of marginal populations. Annu. Rev. Ecol. Syst. 4, 165–187.

Swab, R. M., Regan, H. M., Matthies, D., Becker, U., and Bruun, H. H. (2015). The role of demography, intra-species variation, and species distribution models in species’ projections under climate change. Ecography (Cop.). 38, 221–230. doi:10.1111/ecog.00585.

Therneau, T. M. (2018). coxme: Mixed Effects Cox Models. R package version 2.2–7.

Totland, Ø. (1999). Effects of temperature on performance and phenotypic selection on plant traits in alpine Ranunculus acris. Oecologia 120, 242–251. doi:10.1007/s004420050854.

Totland, Ø., and Alatalo, J. M. (2002). Effects of temperature and date of snowmelt on growth, reproduction, and flowering phenology in the arctic/alpine herb, Ranunculus glacialis. Oecologia 133, 168–175. doi:10.1007/s00442-002-1028-z.

Tutin, T. G., Heywood, V. H., Burges, N. A., Valentine, D. H., Walters, S. M., and Webb, D. A. (1964). Flora Europaea.

Valladares, F., Matesanz, S., Guilhaumon, F., Araújo, M. B., Balaguer, L., Benito-Garzón, M., et al. (2014). The effects of phenotypic plasticity and local adaptation on forecasts of species range shifts under climate change. Ecol. Lett. 17, 1351–1364. doi:10.1111/ele.12348.

Villellas, J., Ehrlen, J., Olesen, J. M., Braza, R., and Garc, B. (2013). Plant performance in central and northern peripheral populations of the widespread Plantago coronopus. Ecography (Cop.). 36, 136–145. doi:10.1111/j.1600-0587.2012.07425.x.

Volis, S. (2011). Adaptive genetic differentiation in a predominantly self-pollinating species analyzed by transplanting into natural environment, crossbreeding and Q ST-F ST test. New Phytol. 192, 237–248. doi:10.1111/j.1469-8137.2011.03799.x.

Wagner, V., Wehrden, H. Von, Wesche, K., Fedulin, A., and Sidorova, T. (2011). Similar performance in central and range-edge populations of a Eurasian steppe grass under different climate and soil pH regimes. Ecography (Cop.). 34, 498–506. doi:10.1111/j.1600-0587.2010.06658.x.

