## Supplementary material including tables, figures caption and more detailed information related to the main text for "Ecological niche models as hypothesis generators of functional genetic differentiation and potential local adaptation in a Mediterranean alpine ecosystem"

***Appendix.***

**Appendix 1: Herbarium records.**

Ten new *Silene ciliata* herbarium specimens deposited in the Real Rardín Botánico de Madrid.

-*Silene ciliata* (Madrid, Rascafría, Pico Dos Hermanas, collector: Javier Morante). Code MA 880770

-*Silene ciliata* (Madrid, Rascafría, morrena lateral Laguna Grande, collector: Mª Luisa Rubio). Code MA 880769

-*Silene ciliata* (Madrid, Rascafría, Pico Peñalara, collector: Carlos Lara). Code MA 880768

-*Silene ciliata* (Madrid, Rascafría, falda pico Najarra, collector: Alfredo García). Code MA 880767

-*Silene ciliata* (Salamanca, Candelario, canchal Negro, collector: Carlos Lara). Code MA 880766

-*Silene ciliata* (Salamanca, La Hoya, Las Cimeras, collector: José Maria Iriondo). Code MA 880765

-*Silene ciliata* (Ávila, Navalperal de Tormes, Los Campanarios, collector: Carlos Lara). Code MA 880764

-*Silene ciliata* (Ávila, Navalperal de Tormes, Alto del Morezón, collector: Javier Morente). Code MA 880763

-*Silene ciliata* (Salamanca, Candelario, Pico El Águila, collector: Mª Luisa Rubio). Code MA 880762

-*Silene ciliata* (Ávila, Navalperal de Tormes, El Servil, collector: Alfredo García). Code MA 880761

**Appendix 2: Thinning distance study**

In the context of this work, it was crucial to control environmental variability loss and thus specific niche signal of the species model sensitivity (too separated presence data not representing the whole environmental range of the species), but also not to neglect sampling biases (too aggregated presence data in some areas).

In order to select the best thinning distance for our study case we explored how much different the variables are between close presence points by comparing their values between close neighbor points at different distances (500m, 1km, 2km). We plotted the relation between environmental differences between presence localities distances and point’s geographical distance following the following steps: (i) select the points with a neighbor less than 500m apart and calculate the distance between them; (ii) extract the environmental values of the variables used in the model (see material and methods) of each point and calculate the absolute value of the difference between them; (iii) plot the relation between geographical distances and the environmental differences. We repeated the procedure for 1km and 2km.

Regarding the geographical distances between presence data that are smaller than 500m, we can see that environmental values are very similar for several groups of data. If we watch geographical distances between presence data between 500m and 1000m, we can see that the environmental differences are not aggregated and that a larger range of environmental differences is represented. Finally, regarding the geographical distances between 1000m and 2000m we can see that as we increase the geographical distances more environmental signal is lost, especially between close presence points and at geographical distances larger than 1500m.

**Appendix 3: Creation and selection of environmental layers**

High resolution climatic layers were needed to develop a fine scale model that would be sensitive to the local steep environmental differences between close areas and thus sensitive to differentiate marginal from optimal habitats. We used fine scale monthly environmental layers from the Atlas Digital Climático de la Península Ibérica with a 200m resolution (ADCPI, Ninyerola et al. (2005), accessed October 2016). ADCPI provides monthly climatic data (minimum, maximum and mean temperature, precipitation and solar radiation) calculated using a temporal coverage of data from 1951-1999. More detailed information can be obtained at: http://opengis.uab.es/wms/iberia/espanol/es_presentacio.htm

Using ENVIREM R package we calculated the 19 widely used WorldClim variables (Hijmans et al., 2005) and potential evapotranspiration (PET). We also calculated monthly snowpack following the methodology proposed by López-Moreno et al. (2007). Based on the previous knowledge on the natural history of the species and its ecosystem (e.g. García-Fernández et al., 2014; Giménez-Benavides et al., 2007; Lara-Romero et al., 2014), exhaustive correlation analysis and experts discussion we selected the variables that best represent the extreme environmental conditions for the species, which made it more likely to capture factors limiting the distribution of the species and to identify areas potentially subjected to strong selective pressures. The final variables used were: minimum temperature of the coldest month (BIO6), precipitation of the driest (BIO14) and the wettest month (BIO13), mean snowpack calculated in thaw months (February, March and April) and mean annual potential evapotranspiration (PET). Environmental variables were constructed and managed using *raster* V2.8-19 and *envirem* V1.4 R packages (Hijmans and van Etten, 2012; Title and Bemmels, 2018). All correlation values between selected variables were lower than 0.7. Correlation analyses were made by the construction of correlation dissimilarity trees/diagrams using *stats* V3.5.3 R packages (Team, 2013).

**Appendix 4: Model environmental space study.**

We also study if the model captured the environmental variability/gradient (retaining species specific niche signal *sensu* Anderson (2012)) inside the study area by plotting the presence data in the environmental space defined by the two first principal components analysis axes made with the environmental variables used in the model. Using this approach we could also study a possible environmental differentiation between mountain ranges inside the Sistema Central by identifying mountain range provenance of each presence data in the same plot and looking for some clustering in the environmental space. The environmental space captured by the whole background sampling was also plotted.

We selected the most important variables used in the model (regarding non-zero lambda values) and made a raster-stack. Then we extracted values of the background points and calculated a PCA (2 axes 97% of the variance); variables were standardized (subtracted mean and divided by standard deviation). After this we extracted the environmental values from the raster-stack of the presence data and projected the values of the presence data in the PCA space created. Then we plotted in the space the background values and the presence points and symbolized them by the suitability values of the model (colors gradient). Also we put labels of the names of 9 potential experimental populations (finally 6 selected for this study) to see if optimal and marginal populations are distinguishable. We also plotted the distribution of the presence data labeled by the mountain ranges (Bejar, Gredos y Guadarrama) to see if there are environmental differences between mountain ranges. All analysis were made using *rgdal* V0.8-16, *rast*er V2.8-19 and *stats* V3.5.3 R packages (Bivand et al., 2014; Hijmans and van Etten, 2012; Team, 2013).

Regarding the figures we can see that the presence data are well distributed along the environmental gradient (figure 4.1A) and that there are not environmental differences between presence data from different mountain ranges (no clustering related with mountain ranges observed in figure 4.1B). Labeled populations selected for the genetic differentiation study seems to be grouped in two different areas inside the environmental space. Populations classified as marginal (Ses, Agi and Naj) have lower habitat suitability values and are located in lower values of the two PCA axes compared to populations classified as optimal (Zon, Neg and Pen).

**Appendix 5: Environmental information from the populations used for the phenology differentiation study and habitat suitability values.**

**Appendix 6: Parameters estimates of the models.**

**References.**

Anderson, R.P., 2012. Harnessing the world ’ s biodiversity data : promise and peril in ecological niche modeling of species distributions. Ann. N. Y. Acad. Sci. 1–15. https://doi.org/10.1111/j.1749-6632.2011.06440.x

Bivand, R., Keitt, T., Rowlingson, B., 2014. rgdal: Bindings for the Geospatial Data Abstraction Library. R package version 0.8-16. URL http//CRAN. R-project. org/package= rgdal.

García-Fernández, A., Escudero, A., Lara-Romero, C., Iriondo, J.M., 2014. Effects of the duration of cold stratification on early life stages of the Mediterranean alpine plant Silene ciliata. Plant Biol. 17, 344–350. https://doi.org/10.1111/plb.12226

Giménez-Benavides, L., Escudero, A., Iriondo, J.M., 2007. Reproductive limits of a late-flowering high-mountain Mediterranean plant along an elevational climate gradient. New Phytol. 173, 367–382. https://doi.org/10.1111/j.1469-8137.2006.01932.x

Hijmans, R.J., Cameron, S.E., Parra, J.L., Jones, P.G., Jarvis, A., 2005. Very high resolution interpolated climate surfaces for global land areas. Int. J. Climatol. A J. R. Meteorol. Soc. 25, 1965–1978.

Hijmans, R.J., van Etten, J., 2012. raster: Geographic analysis and modeling with raster data. R package version 2.0–12.

Lara-Romero, C., Robledo-Arnuncio, J.J., García-Fernández, A., Iriondo, J.M., 2014. Assessing intraspecific variation in effective dispersal along an altitudinal gradient: A test in two Mediterranean high-mountain plants. PLoS One 9. https://doi.org/10.1371/journal.pone.0087189

López-Moreno, J.I., Vicente-Serrano, S.M., Lanjeri, S., 2007. Mapping snowpack distribution over large areas using GIS and interpolation techniques. Clim. Res. 33, 257.

Ninyerola, M., Pons, X., Roure, J.M., 2005. Atlas climático digital de la Península Ibérica: metodología y aplicaciones en bioclimatología y geobotánica. Universitat Autònoma de Barcelona, Departament de Biologia Animal, Biologia Vegetal i Ecologia (Unitat de Botánica).

Team, R.C., 2013. R: A language and environment for statistical computing.

Title, P.O., Bemmels, J.B., 2018. ENVIREM: An expanded set of bioclimatic and topographic variables increases flexibility and improves performance of ecological niche modeling. Ecography (Cop.). 41, 291–307.
