## Supplementary figures and images for "Ecological niche models as hypothesis generators of functional genetic differentiation and potential local adaptation in a Mediterranean alpine ecosystem"

### Supplementary figure A 2.1

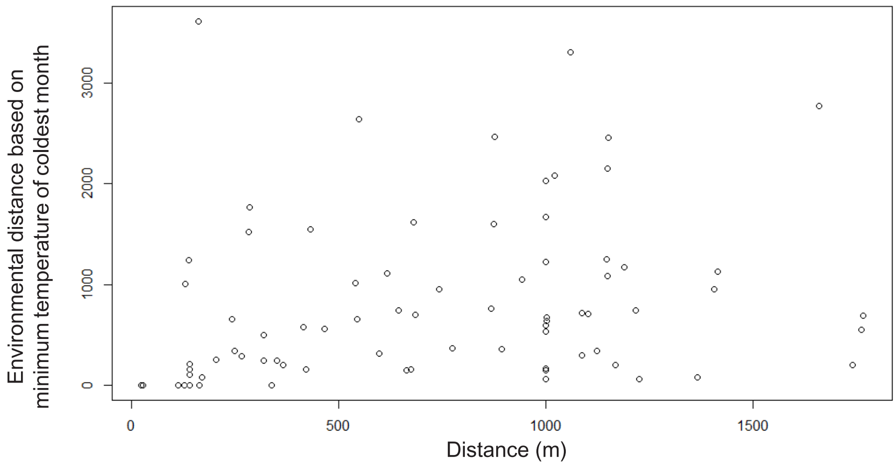

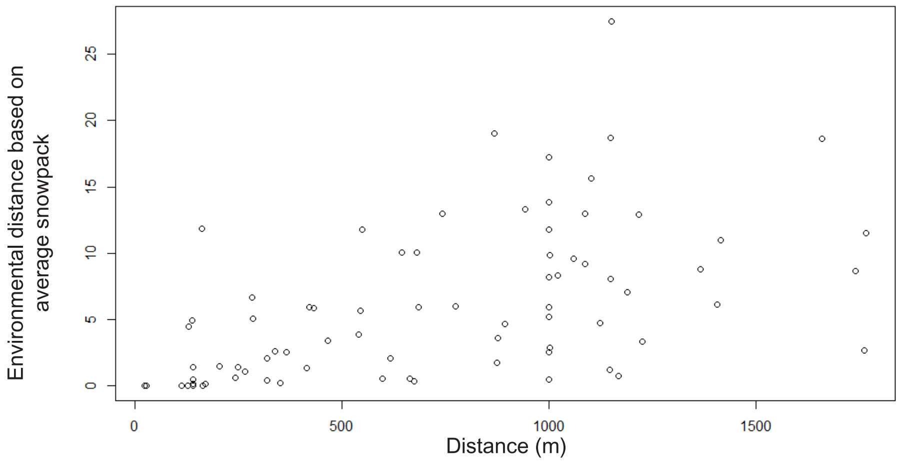

### Supplementary figure A 4.1

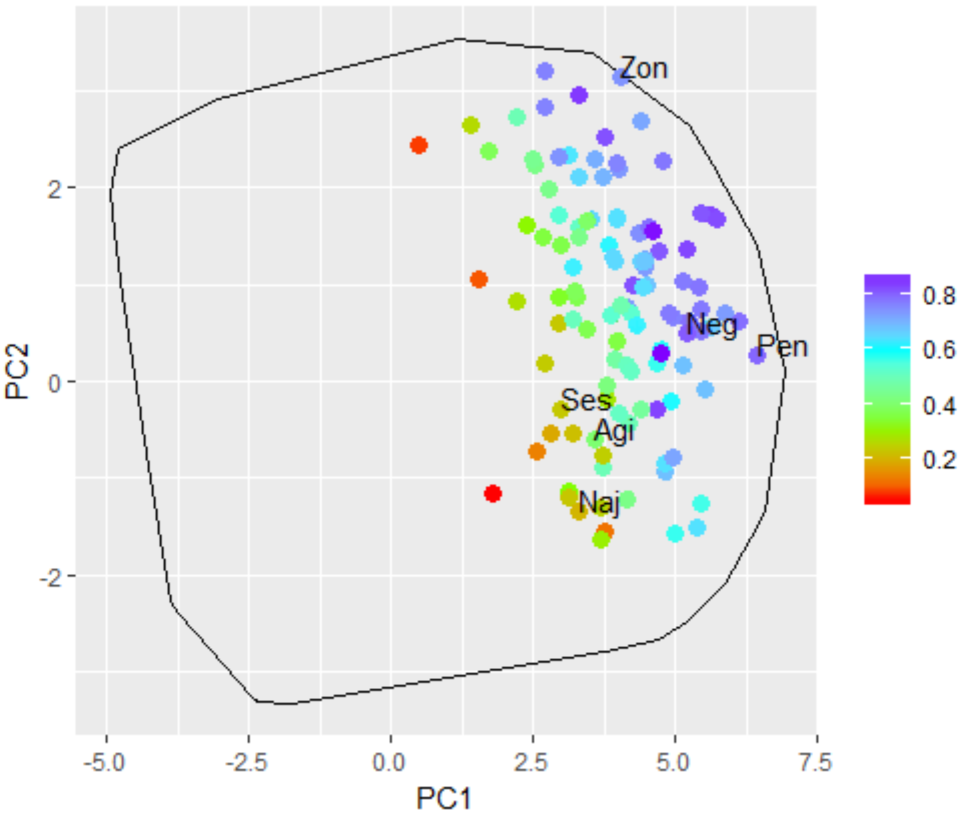

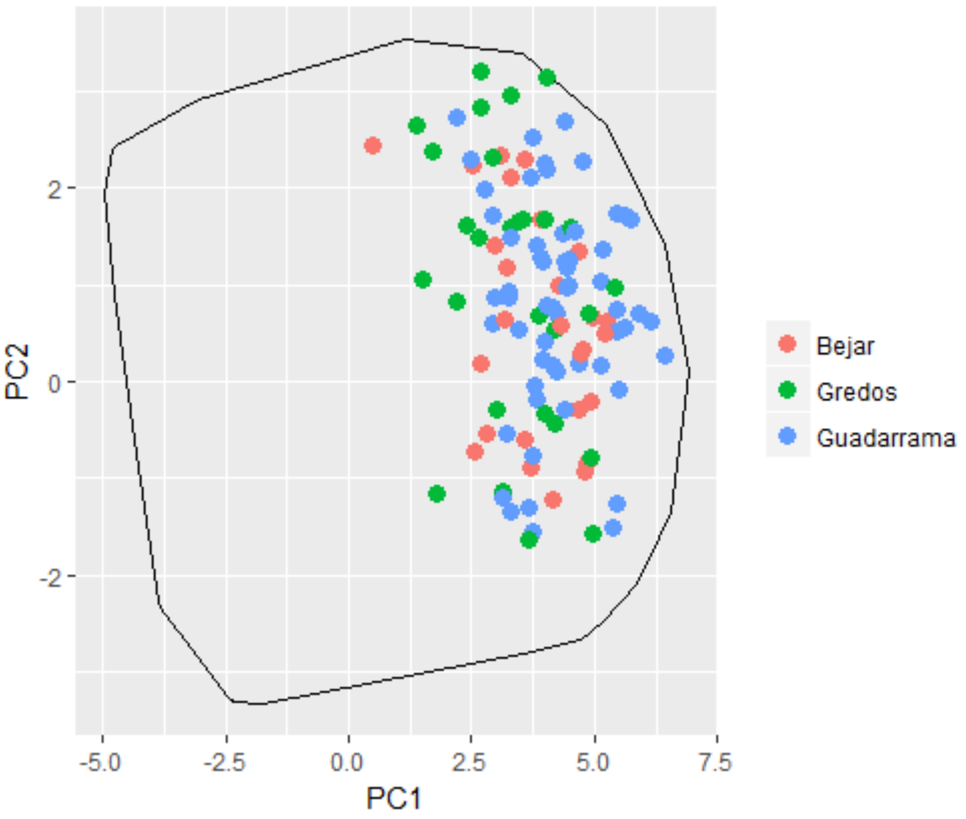
